# Microglia modulate cerebral blood flow and neurovascular coupling through ectonucleotidase CD39

**DOI:** 10.1101/2024.11.05.622122

**Authors:** Zhongxiao Fu, Mallikarjunarao Ganesana, Philip Hwang, Xiao Tan, Melissa Marie Kinkaid, Yu-Yo Sun, Emily Bian, Aden Weybright, Katia Sol-Church, Ukpong B. Eyo, Clare Pridans, Francisco J. Quintana, Simon C. Robson, Pankaj Kumar, B. Jill Venton, Anne Schaefer, Chia-Yi Kuan

## Abstract

Microglia and the border-associated macrophages (BAMs) contribute to the modulation of cerebral blood flow (CBF), but the mechanisms have remained ill-defined. Here, we show that microglia regulate the CBF baseline and upsurges after whisker stimulation or intracisternal magna injection of adenosine triphosphate (ATP). Genetic or pharmacological depletion of microglia reduces the activity-dependent hyperemia but not the cerebrovascular responses to adenosine stimulation. Notably, microglia repopulation corrects these CBF reactivity deficits. The microglial-dependent regulation of CBF requires the ATP-sensing P2ry12 receptor and the ectonucleotidase CD39 that initiates the breakdown of extracellular ATP. Pharmacological inhibition or microglia-specific deletion of CD39 simulates the CBF anomalies detected in microglia-deficient mice and reduces the rise of extracellular adenosine after whisker stimulation. Together, these results suggest that the microglial CD39-initiated conversion of extracellular ATP to adenosine is an important step in neurovascular coupling and the regulation of cerebrovascular reactivity.

## Introduction

Neurovascular coupling, the dynamic link between CBF and neuronal activity, is an important mechanism that safeguards the brain energy demands for cerebral glucose/nutrient and oxygen consumption^1,2^. Among all vasoactive factors, adenosine is considered an important metabolic regulator of CBF^3,4^. In neuronal excitation or cerebral ischemia, ATP is extracellularly released and stepwise converted to adenosine diphosphate (ADP) and adenosine monophosphate (AMP), through the CD39 family of cell-surface ecto-nucleotidases, and then to adenosine by CD73 and tissue-nonspecific alkaline phosphatase (TNAP)^5,6^. The ensuing adenosine elevates CBF primarily through the perivascular A2A receptor (and the A2B receptor with a lower affinity)^5–7^. Besides the adenosine (P1) receptors, the purinergic signaling networks in the central nervous system (CNS) include P2X and P2Y receptors that coordinate the pro-inflammatory and chemotactic functions, respectively^5,8^. Brain microglia and the vasculature endothelial cells express CD39, while deletion of CD39 results in aberrant thromboregulation^9,10^. However, the functions of microglia CD39 with respect to the regulation of microcirculation are yet to be fully characterized^11–15^.

Parenchymal microglia and the CNS border-associated macrophages (BAMs) are brain-resident mononuclear cells. They both exert phagocytosis and other functions to maintain the brain homeostasis^16–19^. Microglia and BAMs have discrete markers, but both have close contacts with cerebral blood vessels to form the juxta-vascular microglia and perivascular macrophages^13,14,20,21^. Microglia detect the extracellular ATP and ADP, mainly through the high-affinity P2Y12 receptor (P2ry12), and either repair or intensify the blood-brain-barrier (BBB) damage in a context– and severity-dependent manner^11,22,23^. A recent study revealed that microglia also provide negative feedback on neuronal activity via CD39-mediated conversion of the activity-dependent ATP into adenosine that in turn acts on the adenosine A1 receptor^12^. Given these findings, we hypothesize that microglia may also modulate neurovascular coupling through the CD39-initiated catalysis of extracellular ATP and ADP, ultimately leading to adenosine.

To test this idea, we compared the impacts of several microglia manipulations: including microglia ablation-and-repopulation through reversible inhibition of the colony-stimulating factor 1 receptor (CSF1R), chemogenetic activation of microglia, and the use of Csf1r^ΔFIRE/ΔFIRE^ (FIRE) mice that are selectively deficient in parenchymal microglia. Readouts included the CBF baseline and the CBF reactivity to whisker stimulation and intra-cisterna magna (ICM) injection of ATP or adenosine^24–26^. We also compared the impacts of microglia depletion on the expression of CD39, CD73, and cyclooxygenase 1 (COX1). Moreover, we compared the effects of microglia-specific deletion of CD39 or CD73 on CBF on activity-dependent surge of extracellular adenosine in the barrel cortex. Our results suggest that the microglia CD39-initiated adenosine synthesis is an important step of neurovascular coupling and the CNS purinergic signaling.

## Results

### Microglia are a new component of the neurovascular unit

To assess the roles of microglia in CBF regulation, we took advantage of the CSF1R inhibition-mediated microglial depletion-and-repopulation after the withdrawal of CSF1R-inhibitors^24^. We applied laser speckle contrast imaging (LSCI) to compare the baseline and whisker stimulation-induced CBF in the same mouse longitudinally: i.e. before the PLX3397 chow treatment (with the baseline microglia density), 9 days after the PLX3397 chow (microglia-ablation), and at 9 days after switching back to normal chow (microglia-repopulation) (Fig. 1a)^24,27,28^. We first confirmed that the microglia density was significantly reduced under PLX3397 chow (∼90% reduction), but near-fully recovered after returning to normal chow for 9 days (Fig. 1b-c and Extended Data Fig. 1a-b). Neither the PLX3397 treatment nor microglia-repopulation altered the GFAP^+^ astrocyte numbers, AQP4 expression (Extended Data Fig. 1c-h), the baseline and post-whisker stimulation cFos^+^ neurons (Extended Data Fig. 1i-k), or the BBB permeability (Extended Data Fig. 2a-b), all in keeping with the previous reports^14,24,29,30^.

**Figure 1.**
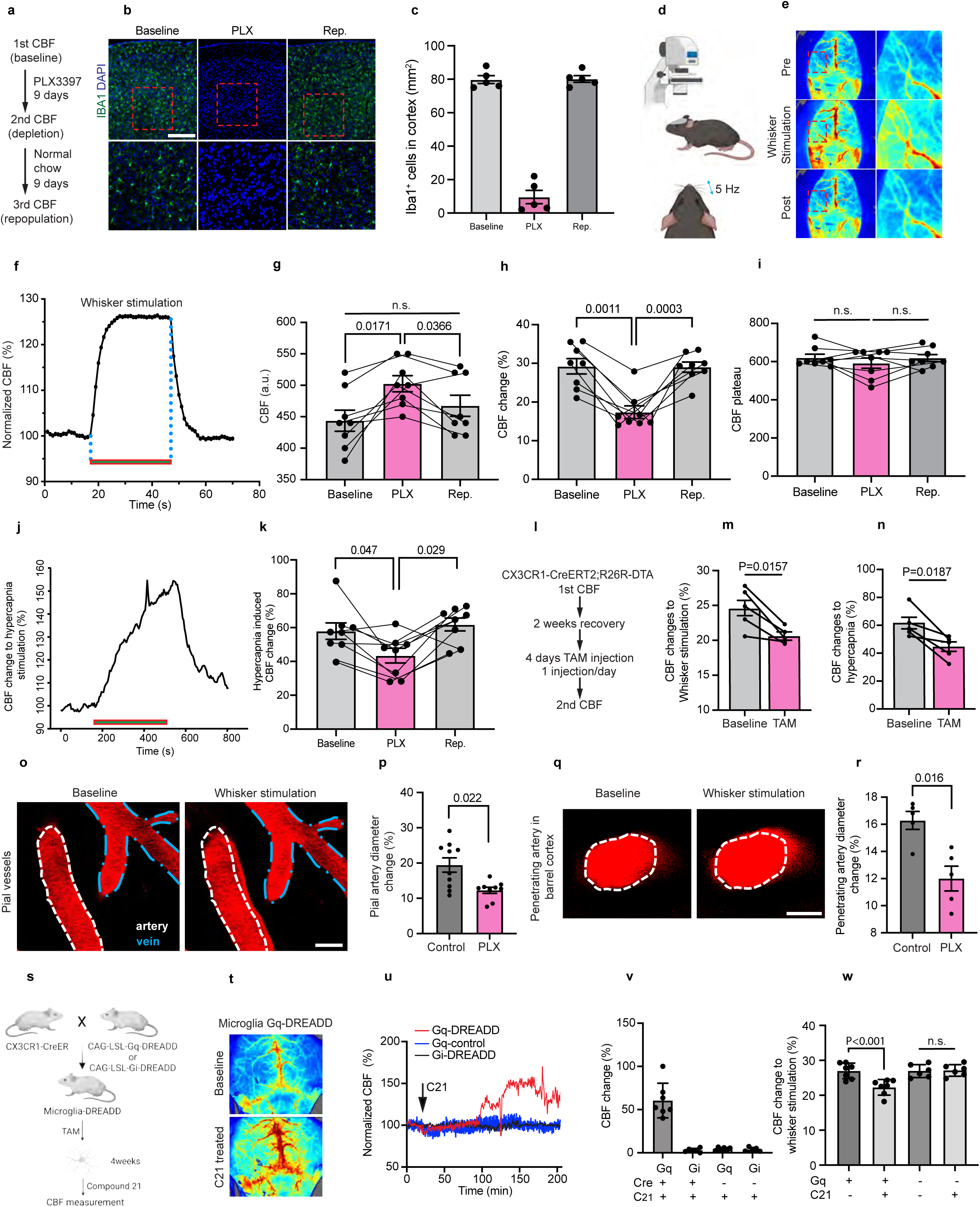
Microglia regulate basal cerebral blood flow and neurovascular coupling. **a,** Schematics for longitudinal comparison of CBF reactivity in the same mouse: the baseline, the microglia-depletion state after PLX3397 chows, and microglia-repopulation after returning to normal chows. **b,** Immunohistochemistry images showing IBA1^+^ cells co-stained with DAPI in baseline, PLX and repopulation conditions. Bar=400μm. Higher magnification images are shown below. Bar=150μm. **c,** Quantification of IBA1^+^ cells in **b**. **d,** Laser speckle contrast imaging was used to measure the CBF responses to whisker stimulation across an intact skull. **e, Left,** Representative images of mouse CBF changes pre-, during and post-whisker stimulation. Red square boxes (enlarged on **Right**), denote the region for CBF quantification in whisker stimulation. **f,** Representative real-time CBF responses to 5Hz whisker stimulation (marked by the bar; 30 sec). **g-i,** Comparison of the basal CBF (**g**), whisker stimulation-induced CBF changes (**h**) and CBF plateau (**i**) at the baseline, PLX-treated, and repopulation states. Each dot indicates an individual mouse. Data are presented as mean ± s.e.m. p-values were determined by one-way ANOVA with Tukey post hoc test. **j,** Representative real-time CBF responses to 8% CO2 hypercapnia stimulation (marked by the bar; 5min). **k,** Comparison of the hypercapnia-induced CBF changes at the baseline, PLX-treated, and repopulation states. Data are presented as mean ± s.e.m. p-values were determined by one-way ANOVA with Tukey post hoc test. **l,** Scheme of CBF evaluations at the baseline (1^st^ CBF) and tamoxifen-induced acute microglia ablation (2^nd^ CBF) in *Cx3cr1-CreERT2;R26R-DTA* mice. **m, n,** CBF changes to whisker stimulation (**m**) and hypercapnia (**n**) at the baseline and after tamoxifen (TAM)-induced acute microglia ablation in the same mouse. Each line tracks the changes in CBF response in the same mouse. p-values were determined by paired *t-*test. **o,** Representative micrographs showing different responses to whisker stimulation in the pial artery (white lines, dilation) and the pial vein (blue lines, no visible dilation) above the contralateral barrel cortex. Bar=30μm. **p,** Comparison of whisker stimulation-induced dilation of a pial artery in the mice under control chow or PLX3397 chow. Each dot indicates an individual mouse. p-values were determined by unpaired *t*-test. Data are presented as mean ± s.e.m. **q,** Representative micrographs showing dilation of the penetrating artery in the barrel cortex after whisker stimulation. Bar=10μm. **r,** Comparison of whisker stimulation-induced dilation of penetrating artery in barrel cortex in the mice under control chow or PLX3397 chow. p-values were determined by unpaired *t* test. Data are presented as mean ± s.e.m. **s,** Schematics for the generation of transgenic mice expressing microglia-specific Gq-DREADD and Gi-DREADD for compound 21-mediated chemogenetic manipulation. **t,** Representative micrographs showing CBF baseline and compound 21 (C21)-induced CBF change in microglia-specific Gq-DREADD expression mice. **u,** Real-time CBF responses after intraperitoneal injection of C21 (arrow) in mice of the labeled genotype. **v,** Summary of microglia Gq or Gi activation-induced CBF changes in different conditions. **w,** Comparison of the CBF changes to whisker stimulation in microglia-Gq mice and WT littermates with or without the C21 injection. ***: p<0.0001 as determined by paired *t*-test.

LSCI showed that 5 Hz whisker stimulation elevated the CBF to a plateau within 10 sec in the contralateral barrel cortex, which declined rapidly after ending the whisker stimulation (Fig. 1d-f). Through serial recording in the same mouse, we found that the PLX3397-treatment elevated the CBF baseline and a blunted response to whisker stimulation without altering the peak level of CBF (Fig. 1g-i). These effects appeared dependent on microglia, since the CBF baseline and CBF reactivity to whisker-stimulation both returned to normal levels after microglia repopulation (Fig. 1g-i). The PLX3397 treatment also blunted the CBF reactivity to hypercapnia (8% CO_2_), but not the peak CBF value (Fig. 1j-k, Extended Data Fig. 2c-d). These results suggest that microglia normally suppress the CBF baseline but support the cerebrovascular reactivity.

Besides CSF1R-inhibition, we also tested the effects of genetic ablation of microglia (Fig. 1l; crossing CX3CR1-CreER and R26R-DTA mice and using tamoxifen to induce the diphtheria toxin A-chain expression^31^). Similar to PLX3397 treatment, the DTA-mediated microglia ablation reduced the CBF response to both whisker stimulation and hypercapnia (Fig. 1m-n). Further, *in-vivo* two-photon microscopy confirmed that whisker-stimulation dilates both pial and penetrating arteries, but not pial veins, in a PLX3397-reducible manner (Fig. 1o-r).

Further, we examined the effects of chemogenetic manipulation of microglia on CBF. We crossed CX3CR1-CreER mice with CAG-LSL-Gq or CAG-LSL-Gi mice to raise the intracellular calcium or decrease cAMP in microglia, respectively, after intraperitoneal injection of compound 21 (C21) (Fig. 1s)^25,32,33^. We found that acute Gq-activation induced prolonged elevation of the CBF, while Gi-activation had no obvious effects (Fig. 1t-v). The C21 treatment also decreased the CBF responses to whisker stimulation in microglia Gq-expressing mice, but not control littermates (Fig. 1w). Together, these results affirm the reports of blunted neurovascular coupling by microglia ablation^15^, and suggest that microglia normally suppress the CBF baseline to further expand the neurovascular coupling reserve.

### Identification of microglia-associated genes for CBF regulation

Neurovascular coupling involves multiple vasoactive effectors and their unique biosynthesis and metabolizing enzymes^34,35^. To investigate the microglial mechanisms on CBF regulation, we used bulk RNA-Seq to compare the cerebral cortical transcriptome in C57BL/6 mice under the control chow, PLX3397 chow, and PLX3397-to-control chow (Fig. 2a). The principal component analysis (PCA) revealed that the control chow (baseline), PLX3397 chow (PLX) and PLX3397-to-control chow (repopulation) samples are segregated into three clusters (Fig. 2b). The heatmap and volcano plot analysis showed far more down-regulated than up-regulated genes in the PLX-treated group compared to the baseline (Fig. 2c). The significantly down-regulated genes include the expected microglia signature markers (e.g. *Tmem119*, *P2ry12*, *Cx3cr1*, *Trem2*). Notably, the expression of these microglial genes recovered after switching back to normal chow (Extended Data Fig. 3a-c), consistent with near-complete repopulation of microglia based on the staining data (Extended Data Fig. 1a, b). Further, the heatmap of RNA-Seq data and RT-qPCR validation revealed a significant decline of CD39 (*Entpd1*) and COX1 (*Ptgs1*) in PLX3397-treated mouse cortex, which recovered in microglia repopulation (Fig. 2d-e). In contrast, the depletion of microglia had minimal effect on the other vasodilator genes, including nitric oxide synthases (NOS, *Nos1*, *Nos2*, *Nos3*) and multiple ATP-sensitive potassium channels (Extended Data Fig. 3d, e).

**Figure 2.**
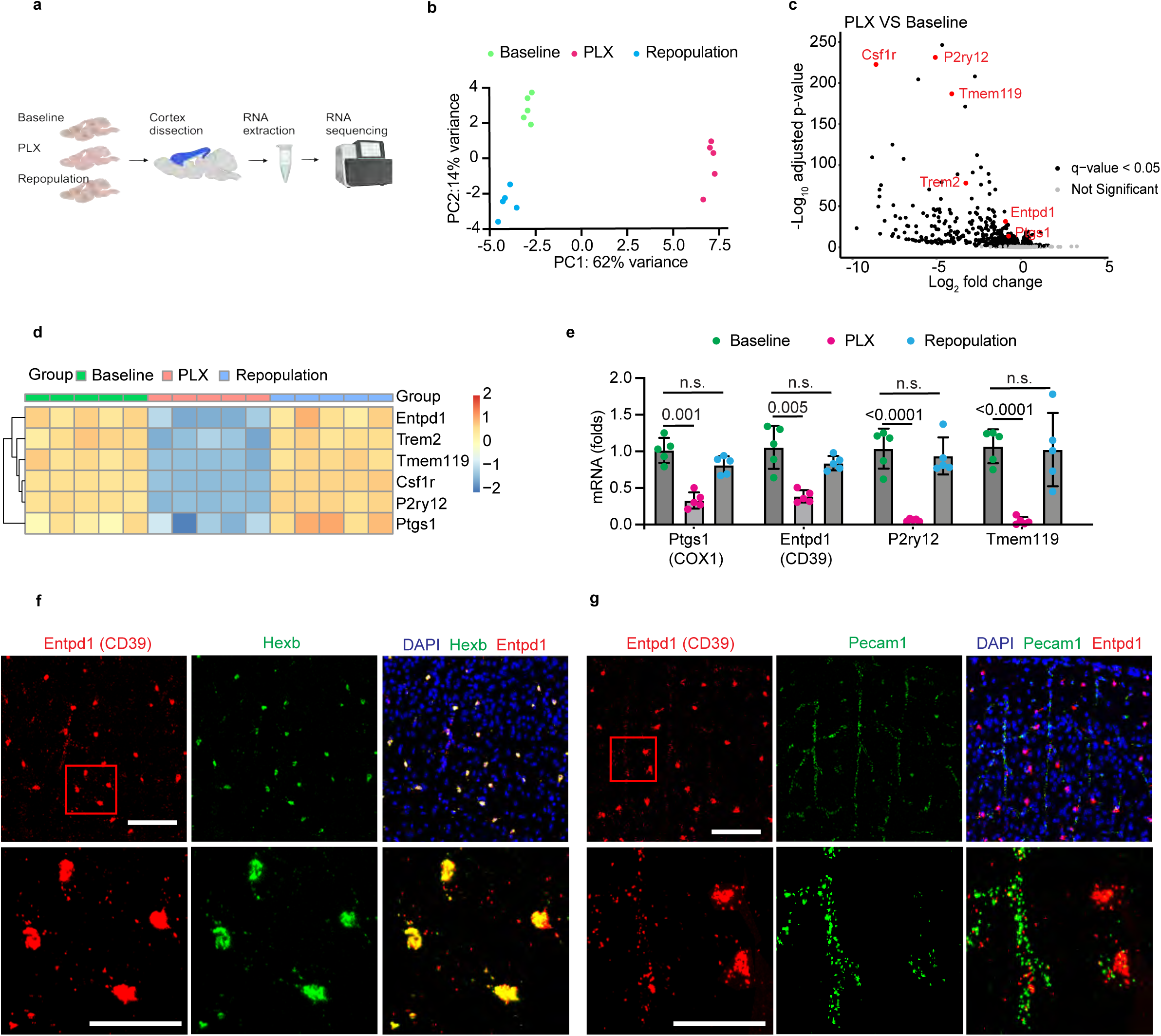
Identification of cerebral blood flow (CBF) regulation-related genes in microglia. **a,** Schematic of bulk RNA sequencing with mice brain cortex tissue from three different treatment groups (baseline, PLX, repopulation). Five mice were included in each group. **b,** Principal component analysis (PCA) plots outlining the similarities between samples. **c,** Volcano plot of the cortex genes showing the magnitude (log2 [fold change]) and probability (−log10 [adjusted p-value]) in the PLX3392-treated group versus the baseline group. Black and gray dots represent significantly changed and non-significantly changed genes, respectively. **d,** Heatmap showing the expression of selected genes in baseline, PLX3397-treated, and repopulated mouse brains based on the row z scores of log2 CPM values. **e,** RT-qPCR verification of the interested genes’ expression in three conditions: baseline, PLX3397-treated and repopulation. Data are mean ± s.e.m. Each dot represents one mouse. p-values were determined by one-way ANOVA with the Tukey post hoc test. **f, g,** Representative images of multi-color RNAscope experiments showing CD39 (Entpd1) expression in microglia (Hexb^+^) (**f**) and endothelial cells (Pecam1^+^) (**g**), respectively. Top, bar=100μm; bottom, bar=50μm.

Previous studies have shown that COX1 modulates CBF and neurovascular coupling^35–37^, but the roles of CD39 in blood flow regulation have not been reported to date. Hence, we used *in-situ* hybridization to examine the *CD39* transcripts and found that they are co-localized with the microglia marker *Hexb* (Fig. 2f)^12,38^ and the endothelial marker *Pecam1* in the penetrating vessels (Fig. 2g), but not in *Aldh1l1^+^* astrocytes (Extended Data Fig. 3g). These results suggest that CD39 is mainly expressed by microglia, and to a lesser degree, in the cerebral vasculature^12,38^. Moreover, the *CD73* transcripts are abundant in the caudate and putamen, but scant in the cerebral cortex (Extended Data Fig. 4a), consistent with little change in the low level of cortical *CD73* transcripts after microglia ablation (Extended Data Fig. 4b). These results also suggests the use of non-CD73 nucleosidases such as TNAP to convert AMP into adenosine in the murine cerebral cortex.

### Differential roles of microglia in ATP-versus-adenosine mediated CBF alterations

Since microglia are the main source of the murine cortical parenchymal CD39, which initiates the conversion of extracellular ATP to adenosine, we conjectured that microglia depletion may have differing impacts on the CBF reactivity to ATP versus adenosine. To test this idea, we used LSCI to compare the CBF after the nano-injector-based intra-cisternal magna (ICM) administration of vasodilator (0.3 μl per min) (Fig 3a). Compared to the cranial window-based CBF study, the ICM-injection method maintains the dura integrity and facilitates longitudinal study of CBF reactivity in the same mouse.

**Figure 3.**
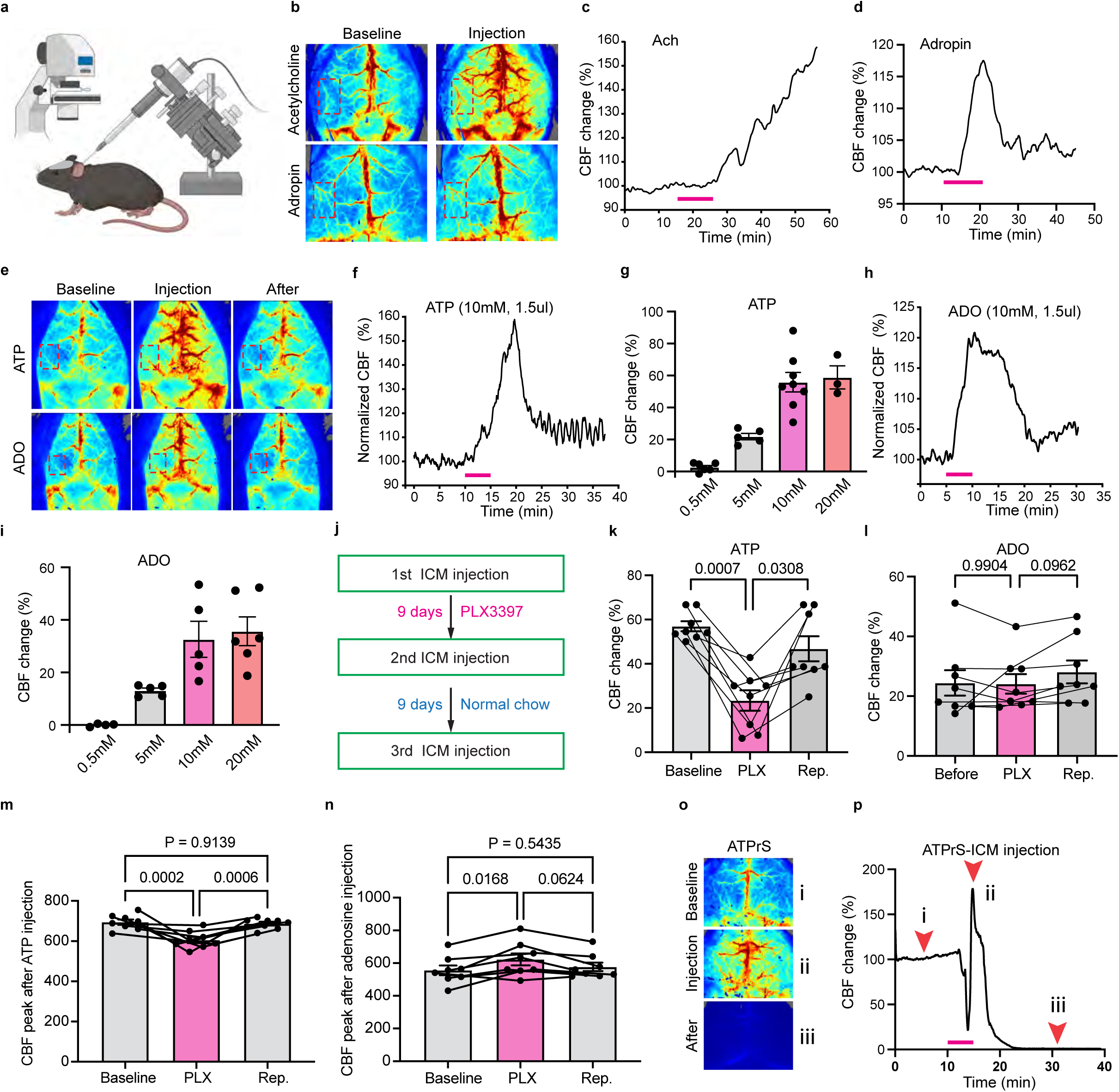
Microglia regulate ATP but not adenosine-induced cerebral blood flow (CBF) increase. **a,** Schematic of combining CBF measurement (using laser speckle contrast imaging) and nano-injector guided intracisternal magna (ICM) injection of vasoactive chemicals. **b,** Representative images of the CBF responses to ICM injection of acetylcholine (**Upper**, 1mM, 3μl) or adropin peptides (**Lower**, 0.1mg/ml, 3μl). **c, d,** Real-time CBF changes to ICM-injection of acetylcholine (Ach, **c**) or adropin (**d**). The bar indicates the 10-minute injection period. **e,** Representative images of the CBF responses to ICM injection of ATP (**Upper,** 1.5μl, 10mM) or adenosine (ADO, **Lower,** 1.5μl, 10mM). **f, g,** Real-time CBF responses to ICM-injection of ATP (**f**) or adenosine (**g**). **h, i,** Does-responses of CBF changes after ICM-injection of ATP (**h**) or adenosine (**i**). The injection volume was 1.5μl in all conditions. **j,** Schematics of sequential measurements of CBF responses in the same mouse at the baseline, after PLX3397-treatment, and at microglia repopulation state. **k, l,** Comparison of the CBF responses to ICM-ATP (10mM, 1.5μl) (**k**) or adenosine (**l**) injection in the same mice at the baseline, PLX-treated, and repopulation state, respectively. Each dot indicates an individual mouse. Data are presented as mean ± s.e.m. p-values were determined by one-way ANOVA with Tukey post hoc test. **m, n,** CBF peak value to ICM-ATP (10mM, 1.5μl) (**m**) or adenosine (**n**) injection in the same mice at the baseline, PLX-treated, and repopulation state, respectively. Each dot indicates an individual mouse. Data are presented as mean ± s.e.m. p-values were determined by one-way ANOVA with Tukey post hoc test. **o, p,** Representative images of the CBF baseline and responses to ICM-injection of ATPγS (10mM, 1.5μl) at the time-points **i**, **ii** and **iii** (indicated by arrowheads in **p**). All ICM-ATPγS-injected mice expired within 20 minutes (n=5).

Using this preparation, we first confirmed the vasodilatory effects of acetylcholine (ACh) and adropin (Fig. 3b-d)^39,40^. Next, we compared the dose-responses of CBF to ATP (Fig. 3e-g) and adenosine (ADO) (Fig. 3e, h, i). Under the same dose for ICM injection (1.5 μl of 10 mM), ATP induced a slower, but greater CBF-upsurge than adenosine (Fig. 3f, h). We then performed serial ICM-CBF assays, first at the baseline, then at 9 days of PLX3397 chow (microglia depletion), and finally at 9 days after returning to control chow (microglia repopulation) (Fig. 3j). We found that microglia depletion significantly blunted the response to ICM-injection of ATP (from 58% to 22% CBF-surge), which recovered after microglia repopulation (returning to 47% of CBF-surge) (Fig. 3k). In contrast, microglia ablation had no effects on CBF responses to ICM-injection of adenosine (Fig. 3l). Consequently, the peak CBF value after ICM-injection of ATP was significantly reduced in microglia ablation despite a higher CBF baseline (Fig. 3m), where the peak CBF after ICM-injection of adenosine was increased (Fig. 3n), due to a higher basal CBF and unaltered responses to adenosine. These results suggest that the microglia-regulated CBF baseline and cerebrovascular reactivity are not coupled to meet a fixed level of plateau CBF.

Finally, we examined the CBF response to ICM-injection of ATPγS, a non-hydrolysable ATP-analogue that cannot be converted to adenosine. We found that ICM-injection of 1.5 μl of 10 mM ATPγS induced a triphasic response of CBF (an initial reduction, a transient surge, invariably followed by vaso-constriction and the animal death) (n=5, Fig. 3o, p). These findings suggest that microglia-initiated ATP breakdown may prevent the lethal effects of non-hydrolysable ATP and attenuate the pro-inflammatory effects of extracellular ATP^4,11^.

### Selective deficiency of parenchymal microglia impairs neurovascular coupling

Because CSF1R-inhibition ablates both parenchymal microglia and BAMs, it remains uncertain whether parenchymal microglia *per se* contribute to the modulation of neurovascular coupling. To address this issue, we utilized the *CSF1R* super-enhancer fms-intronic regulatory element (FIRE) deleted mice (*Csf1r^ΔFIRE/ΔFIRE^*) to assess the CBF-reactivity^26,41^. We first confirmed near-complete absence of Iba1^+^parenchymal microglia in homozygous FIRE mice (and some residual microglia in heterozygous FIRE mice) in the barrel cortex (Fig. 4a, c), but a near-normal density of Lyve1^+^ perivascular and leptomeningeal macrophages (Fig. 4b, d, e; Extended Data Fig. 5a-b). The Lyve1^+^macrophages in choroid plexus were greatly reduced in homozygous, but not heterozygous FIRE mice (Extended Data Fig. 5c, d). Next, we found a significant increase in the CBF baseline and reduction in the reactivity to whisker stimulation, without affecting the peak level of CBF in heterozygous and homozygous FIRE mice, when compared to wildtype littermates (Fig. 4f-h). Further, both heterozygous and homozygous FIRE mice showed diminished CBF-surge after ICM-injection of ATP compared with the wildtype littermates (Fig. 4i-l). These results suggest that the paucity of parenchymal microglia is sufficient to blunt neurovascular coupling and ATP-induced CBF-upsurge. The causes of the initial CBF drop after ICM-injection of ATP in homozygous FIRE mice are uncertain (Fig. 4k), but it may relate to an imbalance between the (absent) microglia and the (remaining) BAM-mediated cerebrovascular functions.

**Figure 4.**
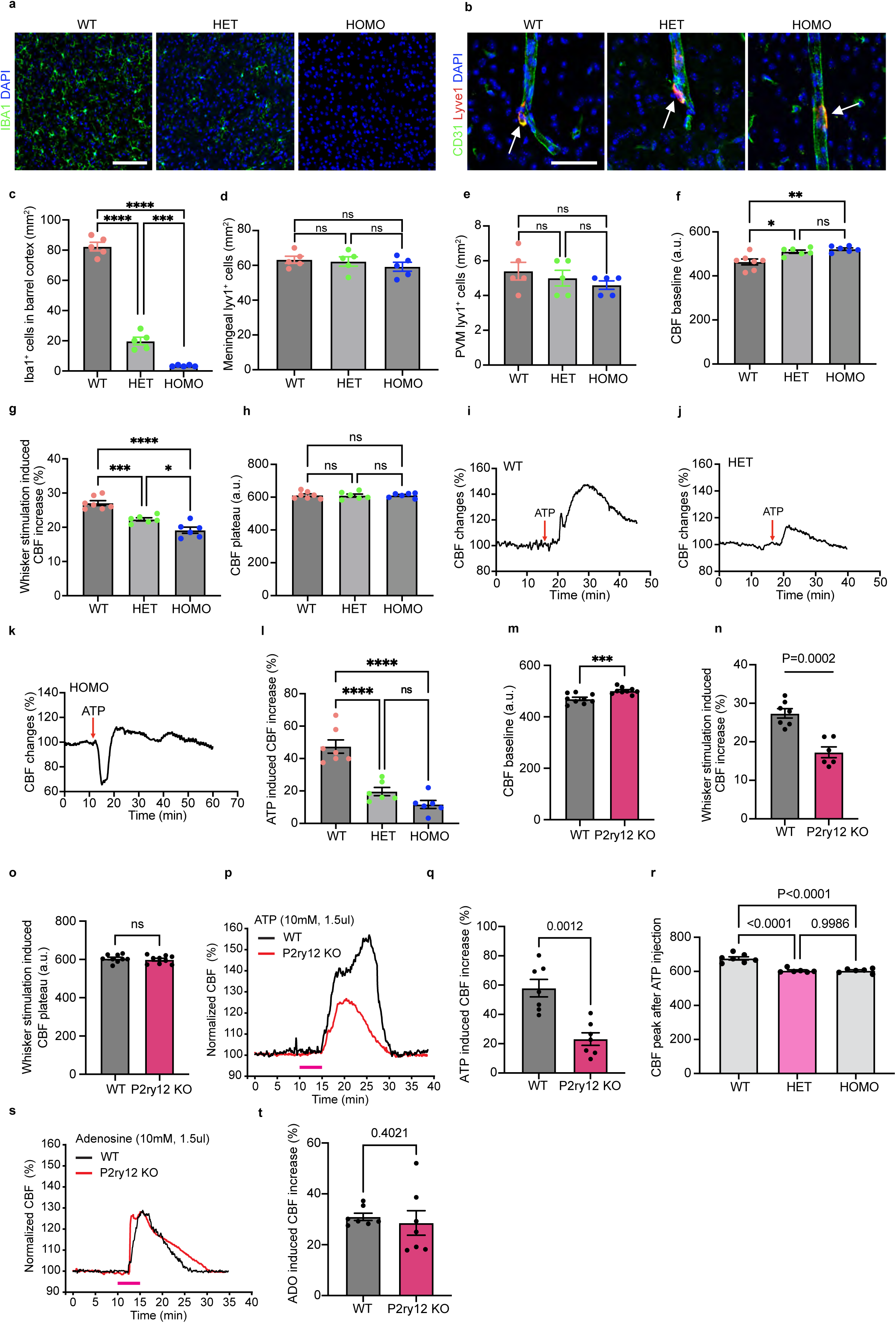
CBF dysregulation in Csf1r^ΔFIRE/ΔFIRE^ and P2RY12 knockout mice. **a,** Representative images showing IBA1^+^ cells (co-stained with DAPI) in barrel cortex of WT, heterozygous, and homozygous Csf1r^ΔFIRE/ΔFIRE^ mice. Bar=200μm. **b,** Representative images showing Lyve1^+^ cells (co-stained with CD31 and DAPI) in barrel cortex of WT, heterozygous (HET), and homozygous (HOMO) Csf1r^ΔFIRE/ΔFIRE^ mice. Bar=50μm. **c-e,** Quantification of IBA1^+^, leptomeningeal Lyve1^+^ and perivascular Lyve1^+^ cell density in WT, HET, and HOMO of Csf1r^ΔFIRE/ΔFIRE^ mice. **f-h,** Quantification of baseline CBF (**f**), whisker stimulation induced CBF changes (**g**) and whisker stimulation induced CBF plateau (**h**) in WT, HET, and HOMO of Csf1r^ΔFIRE/ΔFIRE^ mice. p-values were determined by one-way ANOVA with the Tukey post hoc test. Data are presented as mean ± s.e.m. **i-k,** Representative trances of ATP ICM injection (1.5μl, 10mM) induced CBF changes in WT, HET, and HOMO of Csf1r^ΔFIRE/ΔFIRE^ mice. **l,** Quantification of ATP ICM injection induced CBF changes in WT, HET, and HOMO of Csf1r^ΔFIRE/ΔFIRE^ mice. p-values were determined by one-way ANOVA with the Tukey post hoc test. Data are presented as mean ± s.e.m. **m-o,** Quantification of CBF baseline (**m**), whisker stimulation-induced CBF changes (**n**), and whisker stimulation-induced CBF plateau (**o**). **p**, Example traces showing ATP (1.5μl, 10mM) ICM injection induced CBF changes in WT and P2ry12 knockout mice. **q-r**, Quantification of ATP ICM injection (1.5μl, 10mM) induced CBF changes (**q**) and peak CBF values (**r**) in WT and P2ry12 knockout mice. p-values were determined by unpaired *t*-test in **q**. Data are presented as mean ± s.e.m. p-values were determined by one-way ANOVA with the Tukey post hoc test in **r**. Data are presented as mean ± s.e.m. **s**, Example traces showing adenosine (1.5μl, 10mM) ICM injection induced CBF changes in WT and P2ry12 knockout mice. **t**, Quantification of adenosine ICM injection (1.5μl, 10mM) induced CBF changes in WT and P2ry12 knockout mice. p-values were determined by unpaired *t*-test. Data are presented as mean ± s.e.m.

If microglia need to detect extracellular ATP to initiate its conversion to adenosine for CBF regulation, the microglial P2 purinergic receptors—particularly P2ry12—may also contribute to neurovascular coupling^11,13^. Consistent with this idea, we found an increase in CBF baseline, but diminished CBF reactivity to whisker stimulation in the P2ry12 knockout mice compared with WT mice (Fig. 4m, n), consistent with a past report^15^. Again, the peak CBF after whisker stimulation was unaltered in P2ry12 null mice (Fig. 4o). P2ry12-deficiency also attenuated the CBF responses to ICM-injection of ATP (Fig. 4p-r), but not adenosine (Fig. 4s, t). Of note, P2ry12 is exclusively expressed in parenchymal microglia, but not in BAMs^42,43^. Thus, parenchymal microglia may use P2ry12 to detect the activity-dependent ATP co-transmitter and re-orient the microglial processes to initiate the CD39-mediated ATP-phosphohydrolysis^11,12,44^.

### Microglia CD39 regulates local adenosine concentration in neurovascular coupling

To define the roles of microglial CD39, we assessed the effects of ICM injection of ARL67156, a CD39 inhibitor^12^, and found that it raised the basal CBF in a dose and microglia-dependent manner (Fig. 5a-d and Extended Data Fig. 5f). ICM-injection of ARL67156 (3 μl of 1mM) also blunted the CBF response to whisker stimulation, similar to the effects of microglia depletion (Fig. 5e).

**Figure 5.**
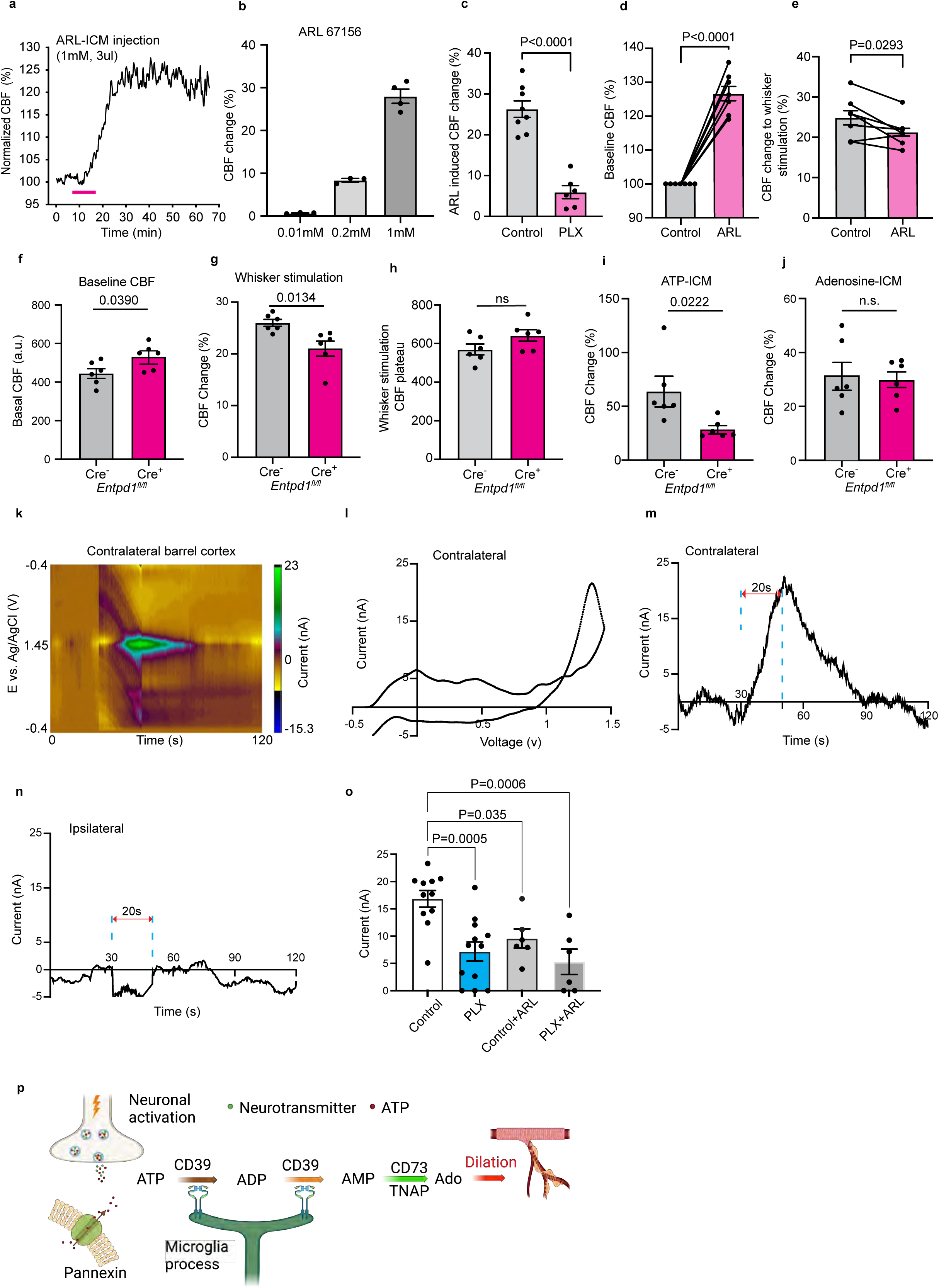
Microglial CD39 regulates basal CBF, neurovascular coupling and whisker stimulation-induced local adenosine concentration increase. **a,** Real-time CBF changes after ICM-injection of ARL67156 (1mM, 3μl). **b,** Does-responses of CBF changes after ICM-injection of ARL67156 (3μl). **c,** The CBF responses to ICM injection of ARL67156 (1mM, 3μl) in control and PLX-treated conditions. Each dot indicates an individual mouse. Data are presented as mean ± s.e.m.. p-values were determined by unpaired *t*-test. **d, e,** Comparison of the CBF baseline (**d**) and the CBF change to whisker stimulation (**e**) in control and at 30min after ICM injection of ARL67156 (1 mM, 3μl). Each line denotes an individual mouse. p-values were determined by unpaired *t* test. **f-i,** Comparison of the CBF baseline (**f**), CBF change to whisker stimulation (**g**), CBF plateau to whisker stimulation (**h**), CBF change to ICM-ATP injection (**i**), and CBF change to ICM-adenosine injection (**j**) in tamoxifen-induced *Cx3cr1-CreER^-^;Entpd1 (CD39)^fl/fl^* and *Cx3cr1-CreER^+^; Entpd1 (CD39)^fl/fl^* mice. Note that microglial CD39-deleted mice showed elevated basal CBF, and reduced CBF change to whisker stimulation and ICM-ATP injection, but no clear changes in the CBF change to ICM-adenosine injection. Each dot indicates one individual mouse. Data are presented as mean ± s.e.m. P-values were determined by unpaired *t*-test. **k-l,** A 3-D color plot of whisker stimulation-induced cyclic voltammogram signals in contralateral barrel cortex (**k**). The 3-D color plot depicts the time on the x-axis, potential on the y-axis, and current in false color. Background subtracted cyclic voltammogram showed the primary oxidation at 1.4 V and the secondary oxidation at 1.0 V (**l**), which are typical for adenosine. **m-o,** Current-vs-time plot shows the whisker stimulation-induced adenosine currents on the contralateral (**m**) and ipsilateral (**n**) barrel cortex. Whisker stimulation was started in the 30s and ended at 50s. Blue dish line marked the 20 seconds stimulation window. **o,** Summary of whisker stimulation-induced adenosine release in different conditions. Each dot indicates one individual mouse. p-values were determined by one-way ANOVA with Tukey post hoc test; * P<0.05, ** P<0.01, *** P<0.0001. Data are presented as mean ± s.e.m. **p,** Schematics of the deduced mechanism in this study by which microglia modulate neurovascular coupling. Upon neuronal excitation such as whisker stimulation, ATP is released from neurons or glia, and undergo microglia CD39-initiated hydrolysis, followed by CD73 or tissue-nonspecific alkaline phosphatase (TNAP)-mediated hydrolysis to form adenosine, leading to vasodilation.

Next, we crossed CX3CR1-CreER and *Entpd1* (CD39)-floxed mice to test the effects of microglial CD39-deletion on CBF. *Cx3cr1^CreERT^*^2^*^/+^;Entpd1^fl/fl^* mice showed near-complete absence of CD39 expression in microglia at 5 weeks after tamoxifen-dosing (Extended Data Fig. 6), when compared to tamoxifen-treated *Entpd1^fl/fl^* mice. These microglial CD39-deleted mice also showed a higher CBF baseline and blunted responses to whisker stimulation and ICM-injection of ATP (Fig. 5f-i), but normal reactivity to ICM-injection of adenosine (Fig. 5j), similar to the PLX3397-treated mice. The CBF plateau under whisker stimulation also remained unaltered in CD39-deleted mice (Fig. 5h). In contrast, the tamoxifen-dosed *Cx3cr1^CreERT^*^2^*^/+^;Nt5e ^fl/fl^* and *Nt5e ^fl/fl^* mice showed normal CBF baseline and reactivity to whisker stimulation and ICM-injection of ATP (Extended Data Fig. 7), in keeping with the scant expression of *Nt5e* (CD73) in the murine cortex (Extended Data Fig. 4). These results implicate microglial CD39, but not CD73, for neurovascular coupling.

Since ATP is an excitatory co-transmitter in most, if not all, synapses in the central and peripheral nervous system^4^, the juxta-synapse and perivascular positioned microglia may engage CD39 to initiate the conversion of extracellular activity-dependent ATP into adenosine to induce physiological hyperemia. To test this idea, we used fast-scan cyclic voltammetry to monitor the response of extracellular adenosine to whisker stimulation in the mouse barrel cortex^45^. Consistent with our hypothesis, whisker stimulation triggered a rapid ascent of extracellular adenosine in the contralateral (Fig. 5k-m), but not ipsilateral barrel cortex (Fig. 5n). Further, microglia ablation and CD39-inhibition by ARL67156 blunted the whisker stimulation-induced extracellular adenosine, without clear synergistic effects (Fig. 5o). These results suggest that microglia CD39 has a critical role in the activity-dependent production of adenosine to promote neurovascular coupling.

## Discussion

Parenchymal microglia have close contacts with both synapses and the cerebral vasculature, but their functional contributions to neurovascular modulation are only coming to light recently^12^. We and others recently reported that dual-depletion of parenchymal microglia and BAMs via CSF1R inhibition blunts the cerebrovascular reactivity to hypercapnia and whisker stimulation^14,15^, but the mechanisms remained unclear. In the present study, we make three new findings regarding the role and mechanisms of microglia in neurovascular modulation and the CNS purinergic signaling.

First, our results ascertain an important role of parenchymal microglia in neurovascular modulation. Since previous studies with CSF1R inhibitors depleted both parenchymal microglia and BAMs, their individual contributions to neurovascular coupling were uncertain^14,15^. Notably, selective ablation of BAMs by intracerebroventricular injection of clodronate liposomes yielded conflicting reports in the effects on neurovascular coupling^15,46,47^. In our study, we used the FIRE mice that lack parenchymal microglia but retain perivascular and leptomeningeal macrophages, which enable us to deduce the functions of parenchymal microglia^26,41^. Our results indicated that the absence of parenchymal microglia is sufficient to diminish CBF responses to ATP– and whisker stimulation (Fig. 4). Given the proximity of microglial processes to both synapses and the cerebral vasculature, we suggest that microglia may transduce neuroexcitation signals to relax pre-capillary arterioles or trigger retrograde hyperpolarization from the capillary to promote activity-dependent hyperemia (i.e. neurovascular coupling)^2^.

Second, our results suggest that adenosine, and specifically, the ecto-nucleotidase CD39, is the critical effector for microglia-mediated neurovascular coupling. The cerebral tissue contains high level of intracellular ATP and adenosine, as well as, a small amount of extracellular ATP and adenosine that has neurovascular modulatory effects^3,4^. ATP has been found to be a co-transmitter in most synapses in the central and the peripheral nervous system where it is released into the extracellular space in an activity-dependent fashion^48^. In addition, ATP can be released through multiple ATP-channels from neuroglia in response to activation, hypoxia, cerebral ischemia, and many other types of brain injury^49–52^. In contrast, adenosine does not appear to be a co-transmitter. Instead, extracellular adenosine arises from the intracellular pool via equilibrative transporters (ENT) or the breakdown of extracellular ATP, first to ADP and AMP through ecto-nucleotidase(s) and then to adenosine via CD73 or soluble TNAP^49^. Extracellular adenosine in turn suppresses neuronal activation or induce vasodilation by the A1– and A2-type adenosine receptors^4,7,12^.

While microglia and the vascular endothelium are known to express CD39^9^, it remains unclear whether CD39 is the principal ecto-nucleotidase to initiate the breakdown of extracellular ATP in cerebral tissue. It is also unclear whether the endothelium-expressed CD39 can compensate for the absence of microglial CD39 to maintain ATP-to-adenosine conversion in the extracellular space in cerebral tissue. Our results demonstrate that depletion of microglia and BAM markedly decreases the transcripts of CD39 in the murine cerebral cortex (Fig. 2e), and that CD39-inhibition or microglia-specific CD39 deletion selectively blunts the CBF responses to whisker stimulation and intra-cisterna magna injection of ATP, but not adenosine (Fig. 5e-j). Further, previous studies showed that whisker stimulation elevates extracellular ATP^50,51^, we add that whisker stimulation induces extracellular adenosine in contralateral barrel cortex in a microglia and CD39-dependent manner (Fig. 5m, o). These results suggest that microglia CD39 is the essential ecto-nucleotidase to initiate the breakdown of activity-dependent extracellular ATP into adenosine for neurovascular coupling in the murine cortex (Fig. 5p).

The microglia CD39-dependent breakdown of extracellular ATP into adenosine provides three entwined benefits (Fig. 5p). First, it prevents the buildup of extracellular ATP, which would allow the influx of Na^+^ and Ca^2+^ through the ionotropic P2X channels to damage neurons and/or activate astrocytes and microglia^53^. Extracellular ATP also induces vasoconstriction after an initial vasodilatory effect^54^, which could have adversarial effects, as suggested in our results using the non-hydrolysable ATPγS (Fig. 3n). These effects of CD39 bioactivity may collectively contribute to the microglia-mediated neuroprotection after stroke^55^. Second, adenosine has depressant effects on cerebral cortical neurons as a negative feedback of neuronal activity, while microglia-depletion or CD39 ablation amplifies neuronal activity and synchronization, leading to seizures^12,56–59^. Third, the microglia CD39-dependent conversion of ATP to adenosine induces physiological hyperemia, as shown in our study, to match the oxygen and glucose/nutrient supply with the neuronal activity. Given these intertwined effects, the microglia CD39 is an integral and important component of purinergic signaling in the brain. Furthermore, our results affirm neurovascular deficits in P2ry12-null mice, presumably due to the absence of P2ry12 in parenchymal microglia^14,15^. Since P2ry12 mediates the chemotactic response to ADP/ATP in microglia^11,13^, this movement may re-orient the microglial processes to initiate the CD39-mediated breakdown of ADP/ATP. Future studies are warranted to test this scenario. Importantly, chemogenetic activation of Gi in microglia simulate the activation of multiple G-protein coupled P2Y receptors besides P2ry12^49^, thus lacking obvious effects on the CBF (Fig 1u-w).

Last but not least, our results suggest that microglia may use several effectors beyond CD39 to exert diverse effects on CBF regulation. Specifically, we showed that chemogenetic induction of calcium in microglial elevates CBF, while the CBF baseline is repressed by microglia (Fig. 1). Since microglia express enzymes for the generation of arachidonic acid metabolites with either vasodilatory or constrictive effects—for example, depletion of microglia markedly decreased the transcripts of COX1 that initiates the metabolism of arachidonic acid (Fig. 2e), microglia may balance the action of various vasoactive effectors to keep an adequate level of basal CBF, while maximizing the cerebrovascular reactivity reserve for responses to urgent needs. Following this conjecture, an imbalance of various microglial vasoactive effectors may cause decreased CBF and glucose metabolism in Alzheimer’s disease and other neurodegeneration disorders^60^. Notably, the expression of microglia CD39 is significantly reduced in the murine models of AD (5xFAD) and amyotrophic lateral sclerosis (SOD1^G93A^)^12,61–63^, which may affect CBF and purinergic signaling.

In conclusion, our results indicate that microglia play a critical role in the CNS purinergic signaling via CD39-dependent catalysis of extracellular ATP to AMP and ultimately to adenosine. Dysregulation of microglial CD39 may lead to impaired cerebral perfusion and cerebrovascular reactivity in stroke, inflammatory diseases and in neurodegeneration.

## Acknowledgements

We thank Ogochukwu Joseph Uweru for his generous sharing of Cx3cr1-GFP mice and Cx3cr1-CreERT2 mice. We thank all the members of the Kuan lab and the members of the Neuroscience Department and Center for Brain Immunology and Glia (BIG) from the University of Virginia for their valuable comments during multiple discussions of this work. We also thank the staff of the University of Virginia Genome Analysis and Technology Core, RRID:SCR_018883, Alyson Prorock and Dr. Yongde Bao, for library preparations and sequencing. *Entpd1^fl/fl^* transgenic mouse was developed with support from National Institutes of Health (HL094400). This work was supported by grants from the National Institutes of Health to C.K. (NS108763, NS127392, NS125788, NS125677), A.S. (AG072489, AG068558, NS106721, MH118329, DA047233, ERC-951515 Micro-COPS), B.J.V. (NS121014), S.C.R. (HL094400, HL167511, AG065923) and P.H. (F31AG074652).

## Methods

### Mouse strains and housing

Wild-type mice (C57BL/6J background) were purchased from the Jackson Laboratory. All mice were maintained in the animal facility for at least one week before to the start of any experiment. The following strains were used: C57BL/6J (JAX 000664), B6N.129-Tg(CAG-CHRM3*,-mCitrine)1Ute/J (CAG-LSL-Gq-DREADD, JAX 026220), B6N.129-Gt(ROSA)26Sor^tm1(CAG-CHRM4*,-mCitrine)Ute/J^ (R26-LSL-Gi-DREADD, JAX 026219), B6.129P2(C)-Cx3cr1^tm2.1(cre/ERT2)Jung/J^ (JAX 020940). P2RY12^−/−^ mice were donated by Dr. Ukpong B. Eyo. Csf1r^ΔFIRE/ΔFIRE^ mice were donated by Dr. Clare Pridans. Csf1r^ΔFIRE/ΔFIRE^ mice used for experiments were crossed with C57BL/6 mice for two generations after importation. All mice were housed under standard 12 h light:dark cycle conditions (lights on at 7:00am) in rooms with controlled temperature and humidity. They were given standard rodent chow and sterilized tap water *ad libitum* unless stated otherwise. Male mice at 10 to 20 weeks of age were used for experiments unless stated otherwise. All experiments were approved by the Institutional Animal Care and Use Committee of the University of Virginia.

To achieve conditional microglia-specific ablation of CD39 or CD73, conditional Cd39^fl/fl^ mice or CD73^fl/fl^ mice^1–3^ were bred to knock in *Cx3cr1^tm2.1(Cre/ERT2)Litt)^*mice (Jackson Laboratory, stock number 021160). All mice used for experiments were backcrossed to the C57Bl/6J background for ≥5 generations. If not otherwise specified, Cre-negative littermate controls were treated with tamoxifen and used as controls. Unless otherwise specified, male and female mice were used for all experiments. Routine genotyping was performed by tail biopsy and PCR as previously described^2^.

### Tamoxifen induction for CD39 KO

Tamoxifen (Sigma-Aldrich) was dissolved in corn oil at a concentration of 20mg/ml. Mice were gavaged with tamoxifen dissolved in corn oil at the concentration of 100 mg/kg body weight, every other day for 6 days. For microglia inducible Cre expression-related experiments, mice were used for experiments four to five weeks after tamoxifen injection.

### Laser speckle contrast imaging

A 2-dimensional laser speckle contrast imaging system following the manufacturer’s instructions (MoorFLPI-2, Moor Instruments) was used to evaluate the CBF dynamics^4,5^. Briefly, mice were anesthetized with urethane (750 mg/kg) and chloralose (50 mg/kg)^6,7^ and placed in the prone position with their skulls exposed but unopened. Mice were put on heating pad and body temperature was continuously monitored during the whole experiment. CBF signal was measured in both cerebral hemispheres. For whisker stimulation-induced real-time CBF dynamics, CBF signaling was recorded at 1Hz for at least 3min stable baseline before 30sec whisker stimulation was applied and recording continued for at least 1min after stimulation. Each mouse was recorded for three rounds of whisker stimulation. The interval between whisker stimulation is 5 minutes. For hypercapnia-induced CBF changes recording, mice were first recorded for 8 to 10min stable baseline under normal air composition and then changed to hypercapnia (8% CO2) condition for 5min, followed by normal air condition. CBF was analyzed by the MoorFLPI software and is shown as arbitrary units in a 16-color palette.

### Intra-cisterna magna injection

Mice were anesthetized with urethane (750 mg/kg) and chloralose (50 mg/kg). Mice were put on a heating pad and body temperature was continuously monitored during the whole experiment. The skin of the neck is shaved and sterilized with iodine. The head of the mouse was gently secured in a stereotaxic frame. A skin incision was made, and the muscle layers were retracted to expose the cisterna magna. Using a WPI nanoliter 2020 injector coupled with pre-pulled glass pipettes to inject the drug into the cisterna magna compartment. All drugs were dissolved in fresh artificial cerebrospinal fluid (119 mM NaCl, 26.2 mM NaHCO_3_, 2.5 mM KCl, 1 mM NaH_2_PO4, 1.3 mM MgCl_2_, 2.5mM CaCl_2_, 10 mM glucose) for ICM injection. The skin was sutured, and the mice were allowed to awake up before returning to their original cage.

### Acute and chronic window implantation

Mice were implanted with a cranial window as previously described^8^. Briefly, during surgery, mice were anesthetized with isoflurane (5% for induction; 1.5% for maintenance) and placed on a heating pad. Using a dental drill, a circular craniotomy of ∼3 mm diameter was drilled at 0.8mm posterior to bregma and 3mm lateral and the center was barrel cortex. A 70% ethanol-sterilized 3 mm glass coverslip was placed above the craniotomy. A light-curing dental cement (Tetric EvoFlow) was applied and cured with a Kerr Demi Ultra LED Curing Light (DentalHealth Products). iBond Total Etch glue (Heraeus) was applied to the rest of the skull, except for the region with the window. This was also cured with the LED Curing Light. The light-curing dental glue was used to attach a custom-made head bar onto the other side of the skull from which the craniotomy was performed. For brain vascular imaging, imaging windows were monitored immediately after the surgery.

### Two photon imaging

The cranial window was prepared above the barrel cortex in wildtype mice treated with either control food or PLX3397 food for 9days. Rhodamine B dye (2 mg/mL) was injected intraperitoneally or subcutaneously to label the vasculature. Mice were kept on heating pad during the whole imaging period. For measuring vascular responses to whisker stimulation, after obtaining 2 min at baseline, a 30-second episode of whisker stimulation was applied and followed by 1min post-stimulation period imaging. Imaging was conducted using a Leica SP8 Multiphoton microscope with a coherent laser. A wavelength of 900 nm was optimal for imaging blood vessel dye using a 25 × 0.9 NA objective. Leica Application Suite X and ImageJ were used for image analysis.

### Mechanical whisker stimulation

Mechanical whisker stimulation was performed electromechanically driven by an electric motor (stereotactic neurosurgery burr 78001, RWD) which connected to a cotton swap. Whole device was fixed to stereotaxy. For electromechanically controlled stimulation (5 Hz), whiskers were stimulated for 30 seconds, repeated four to five times at 5-minute intervals. Stimulation-evoked CBF responses in the contralateral barrel cortex were recorded. All coupling experiments performed were time-matched from the time of anesthetic injection to ensure comparable results across different experiments.

### Chemogenetics manipulation of microglia activity and CBF measurement

R26-LSL-Gq-DREADD or R26-LSL-Gq-DREADD were crossed with Cx3cr1^CreErt2/+^(Jung) mice to produce heterozygous off-spring expressing one copy of the chemogenetic Gq-receptor allele or Gi-receptor allele specifically in microglial cells. Mice were injected with tamoxifen (20mg/ml) daily for five consecutive days to induce Cre expression in microglia. 4weeks later, mice were used for chemogenetics experiments. For Gq-DREADD or Gi-DREADD activation, DREADD agonist 21 (Compound 21, Hello Bio) dihydrochloride (0.5mg/kg) was injected intraperitoneally. CBF was monitored for at least 10min as a baseline before compound 21 i.p. injection and continued during whole experiments.

### Tamoxifen-induced gene expression

Tamoxifen (Sigma-Aldrich) was dissolved in corn oil at a concentration of 20mg/ml. Mice were injected with tamoxifen at the concentration of 3mg/20g body weight for 4 to 5 consecutive days. For microglia inducible Cre expression-related experiments, mice were only used for experiments four to five weeks after tamoxifen injection.

### Tissue preparation and RNA purification

To prepare tissues for bulk RNA sequencing, three groups of mice were included, the baseline group, the PLX3397 treated group, and the microglia repopulation group. Each group included five mice. Baseline group mice were treated with control food. PLX group mice were treated with PLX3397 food for 9 days. Microglia repopulation group mice were first treated with PLX3397 food for 9 days and then put back to control food for 9 days. Mice were given a lethal dose of anesthesia by intraperitoneal (i.p.) injection of euthasol (10% v/v in saline). They were transcardially perfused with ice-cold PBS. Whole forebrain RNA was purified as previously described^9^.

### Library Preparation and Sequencing

Samples were processed by the UVA Genome Analysis and Technology Core, RRID:SCR_018883 using Standard Operating Procedures. Total RNA quality was checked using the Agilent Tape Station 4200. RNAseq libraries were prepared using the NEBNext Ultra II Directional RNA Library Prep Kit for Illumina, according to the manufacturer’s instructions. Libraries were checked for quality, size, and concentration using the Agilent Tape Station 4200 High Sensitivity D5000 kit and Qubit 3.0 (ThermoFisher Scientific) dsDNA HS Assay Kit. Libraries were pooled at equimolar concentrations and sequenced.

### Bulk RNAseq analysis

On average we received 30 million paired ends for each of the replicates. RNAseq libraries were checked for their quality using the fastqc program http://www.bioinformatics.babraham.ac.uk/projects/fastqc/. The results from fastqc were aggregated using multiqc software^10^. Reads were mapped with the “splice aware” aligner ‘STAR’, to the transcriptome and genome of mm10 genome build^11^. The HTseq^12^ software will be used to count aligned reads that map onto each gene. The count table was imported to R to perform differential gene expression analysis using the DESeq2 package^13^. Low-expressed genes (genes expressed only in a few replicates and had low counts) were excluded from the analysis before identifying differentially expressed genes. Data normalization, dispersion estimates, and model fitting (negative binomial) was carried out with the DESeq function. The log-transformed, normalized gene expression of 500 most variable genes will be used to perform an unsupervised principal component analysis. The differentially expressed genes were ranked based on the log2fold change and FDR-corrected p-values. The ranked file was used to perform pathway analysis using GSEA software^14^. The enriched pathways were selected based on enrichment scores as well as normalized enrichment scores. The Enrichment Analysis for Gene Ontology was performed by topGO. The heatmaps were generated using the pheatmap function in R.

### Tissue collection and processing

Mice were given a lethal dose of anesthesia by intraperitoneal (i.p.) injection of euthasol (10% v/v in saline). They were transcardially perfused with ice-cold PBS with heparin (10 U ml^-1^) followed by 4% paraformaldehyde (PFA). Brains were dissected and kept in 4% PFA overnight at 4°C. The fixed brains were washed with PBS, cryoprotected in 30% sucrose for 48 h at 4°C, and frozen in Tissue-Plus OCT compound (Thermo Scientific). Frozen brains were sliced into 40 μm-thick free-floating coronal sections using a cryostat (Leica). Brain sections were stored in PBS with 0.02% azide at 4°C until further use.

### Immunohistochemistry

Mice were given a lethal dose of anesthesia by intraperitoneal (i.p.) injection of euthasol (10% v/v in saline). They were transcardially perfused with ice-cold PBS with heparin (10 U ml^-1^) followed by 4% paraformaldehyde (PFA). Brains were dissected and kept in 4% PFA overnight at 4°C. The fixed brains were washed with PBS, cryoprotected in 30% sucrose at 4°C until sunk to the bottom of the vial, and frozen in Tissue-Plus OCT compound (Thermo Scientific). Frozen brains were sliced into 20 μm-thick free-floating coronal sections using a cryostat (Leica). Brain sections were blocked with 1% bovine serum albumin (BSA), 2% normal serum (either goat or chicken), and 0.2% Triton X-100in PBS for 1 h at room temperature (RT). Sections were incubated with primary antibodies overnight at 4°C in a moist chamber. The following primary antibodies were used: Rabbit anti-Iba1 (No. 019-19741, Lot No. 4987481428584, FUJIFILM, 1/500), Rabbit anti-cFos (Cat. No. 226 008, Synaptic Systems, 1/500), Mouse anti-NeuN (Cat. No. MAB377, Millipore Sigma, 1/500), Chicken anti-GFAP (Cat. No. ab4674, Abcam, 1/500), Rabbit anti-AQP4 (Cat. No. AB3594, Millipore Sigma, 1/500), Mouse anti-Lyve1 (Cat. No. 50-0443-82, Invitrogen, 1/500), Rat anti-CD31 (Cat. No. 5537, Biosciences, 1/500). Secondary antibodies were incubated for 1.5h at room temperature. The following secondary antibodies were used: goat anti-rabbit Alexa Fluor 488 (Cat. No. A11009, Fisher Scientific, 1/500), goat anti-mouse AlexaFluor 647 (Cat. No. 405322, Biolegend, 1/500), goat anti-Chicken Alexa Fluor 488 (Cat. No. A-11039, ThermoFisher Scientific, 1/500). Sections were washed two times for 15 min at room temperature (RT) with PBS. The tissues were mounted with Fluoro-Gel II with DAPI (Electron Microscopy Sciences, Lot#220909).

### Single-molecule multiplex fluorescent in situ hybridization with RNAscope

Euthanized mice were perfused with cold PBS containing heparin (10 U ml^-1^). Within 5 min, brains were embedded in Tissue-Plus OCT compound and were immediately frozen on dry ice. Coronal slices of 16 μm thickness were cut, mounted onto Superfrost plus slides and pretreated according to the manufacturer’s instructions for fresh frozen samples (RNAscope® Multiplex Fluorescent Reagent Kit v2 Assay). Briefly, sections were fixed in pre-chilled 10% neutral buffered formalin (Fisher Scientific) for 30 min at 4°C and were dehydrated using an ethanol series (50%, 70%, and 100% ethanol). The sections were pretreated with hydrogen peroxide for 10 min at RT and incubated with protease IV for 30 min at RT, provided in the kit. The hybridization and amplification steps were performed using Entpd1 (Mm-Entpd1-C3), Nt5e (Mm-Nt5e-C2), Hexb (Mm-Hexb-C1), Aldh1l1 (Mm-Aldh1l1-C1), Pecam1 (Mm-Pecam1-C1) RNAscope probes according to manufacturer’s instructions. Detection of Entpd1 (or Nt5e) on microglia, astrocytes or endothelial cells was achieved using TSA Vivid fluorophore 520 (1:1500, ACD) and TSA Vivid fluorophore 650 (1:1500, ACD). Nuclei were stained with DAPI at RT for 30 s and sections were mounted with Fluoro-Gel II with DAPI (Electron Microscopy Sciences, Lot#220909) and covered with coverslips.

### RNA extraction and quantitative real-time PCR

The total RNA of the control or PLX 3397 brain samples was extracted using the TRIzol reagent (Thermo Fisher Scientific); 3 µg RNA was used for cDNA synthesis using High-Capacity cDNA Reverse Transcription Kit (Applied Biosystems) according to the manufacturer’s instructions. qRT-PCR was performed using the Bio-Rad CFX 96 system (C1000 Thermal Cycler) and detected by SYBR Green master mix (Bio-Rad). Primer sequences were used for real time PCR:

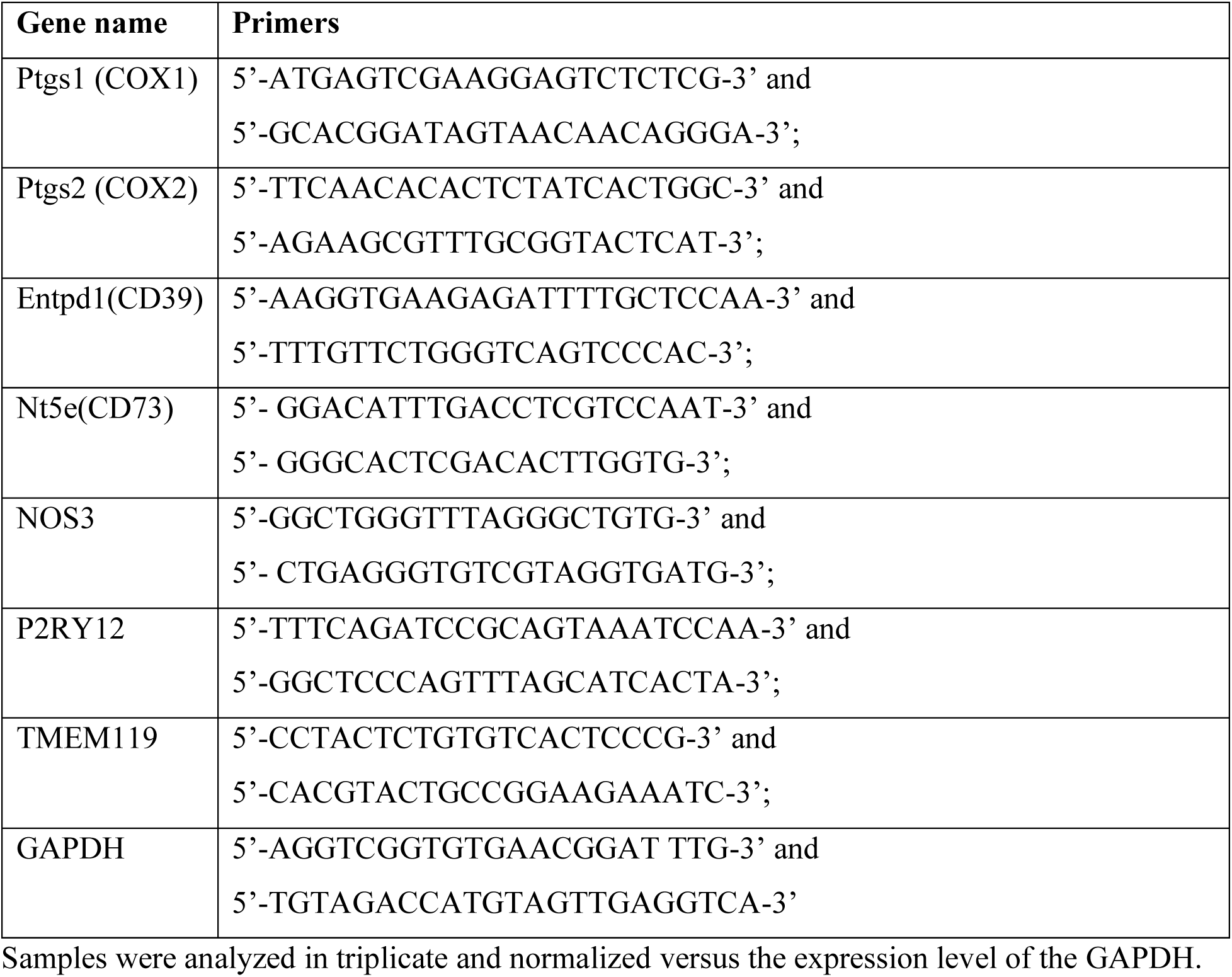

### Fast-scan cyclic voltammetry and electrochemical detection of whisker stimulated adenosine

Carbon-fiber micro-electrodes (CFMEs) were fabricated by inserting a single carbon fiber (T-650, Cytec Engineering Materials) of 7 µm in diameter into a glass capillary (0.68 mm ID × 1.2 mm OD) and pulled using a vertical pipette puller (model PE-21) into two electrodes^15^. The exposed carbon fiber was cut to 125–150 µm with a scalpel. An electrical connection was made by backfilling the capillary with 1 M KCl. The silver-silver chloride reference electrodes were made in-house by electrodepositing chloride onto a silver wire (Acros Organics).

Fast-scan cyclic voltammetry (FSCV) was used to monitor continuous adenosine release in real-time on a sub second time scale in vivo in anaesthetized animals^16,17^. Whisker-stimulated adenosine release was collected through computer-controlled HDCV software (University of North Carolina) using a Dagan Chem Clamp potentiostat (Dagan Corporation). The applied waveform was from –0.40 V to 1.45 V and back at 400 V/s vs Ag/AgCl reference and was repeated for every 100 ms. As FSCV produces a large background current, data were background subtracted (10 cyclic voltammograms averaged) to remove nonfaradaic currents. Electrodes were post calibrated with 1.0 µM adenosine in PBS solution, immediately following animal experiments and the average of triplicate current responses was used to estimate the whisker-stimulated adenosine concentrations in vivo.

### PLX3397 treatment

For microglial depletion studies, mice were fed with chow containing a final dose of 660 mg/kg PLX3397, a CSF1R inhibitor widely used to eliminate microglia from the brain^8^. For microglial repopulation studies, the mice were switched from Plexxikon chow (formulated with Research Diets, New Jersey) back to control chow to allow for microglial repopulation^8^.

## Statistical Analysis

GraphPad Prism version 9, OriginPro 7.5, and MATLAB R2017b were used for statistical analysis. All data were presented as mean ± SEM. N indicates the number of animals unless otherwise specified. For comparisons between two groups or more, data were analyzed by One-Way ANOVA followed by the post-hoc Tukey-Kramer test. For comparisons of distribution data between two-groups, the data were analyzed by unpaired t test.

## Data availability

Raw data for mouse bulk RNA-sequencing have been deposited in the Gene Expression Omnibus, and are available at the following accession numbers: GSE245107. All other data are available from the corresponding author on request.

## References

1 Lassen, N. A. Cerebral blood flow and oxygen consumption in man. Physiological reviews 39, 183–238 (1959).

2 Schaeffer, S. & Iadecola, C. Revisiting the neurovascular unit. Nature Neuroscience 24, 1198–1209 (2021).

3 Berne, R. M., Winn, H. R. & Rubio, R. The local regulation of cerebral blood flow. Progress in cardiovascular diseases 24, 243–260 (1981).

4 Abbracchio, M. P., Burnstock, G., Verkhratsky, A. & Zimmermann, H. Purinergic signalling in the nervous system: an overview. Trends in neurosciences 32, 19–29 (2009).

5 Burnstock, G. Purinergic signaling in the cardiovascular system. Circulation research 120, 207–228 (2017).

6 Taruno, A. ATP release channels. International journal of molecular sciences 19, 808 (2018).

7 Kusano, Y. et al. Role of adenosine A2 receptors in regulation of cerebral blood flow during induced hypotension. Journal of Cerebral Blood Flow & Metabolism 30, 808–815 (2010).

8 Di Virgilio, F., Sarti, A. C. & Coutinho-Silva, R. Purinergic signaling, DAMPs, and inflammation. American Journal of Physiology-Cell Physiology 318, C832–C835 (2020).

9 Braun, N. et al. Assignment of ecto-nucleoside triphosphate diphosphohydrolase-1/cd39 expression to microglia and vasculature of the brain. European Journal of Neuroscience 12, 4357–4366 (2000).

10 Enjyoji, K. et al. Targeted disruption of cd39/ATP diphosphohydrolase results in disordered hemostasis and thromboregulation. Nature medicine 5, 1010–1017 (1999).

11 Haynes, S. E. et al. The P2Y12 receptor regulates microglial activation by extracellular nucleotides. Nature Neuroscience 9, 1512–1519 (2006).

12 Badimon, A. et al. Negative feedback control of neuronal activity by microglia. Nature 586, 417–423 (2020).

13 Cserép, C. et al. Microglia monitor and protect neuronal function through specialized somatic purinergic junctions. Science 367, 528–537 (2020).

14 Bisht, K. et al. Capillary-associated microglia regulate vascular structure and function through PANX1-P2RY12 coupling in mice. Nature communications 12, 5289 (2021).

15 Császár, E. et al. Microglia modulate blood flow, neurovascular coupling, and hypoperfusion via purinergic actions. Journal of Experimental Medicine 219 (2022).

16 Li, Q. & Barres, B. A. Microglia and macrophages in brain homeostasis and disease. Nature Reviews Immunology 18, 225–242 (2018).

17 Paolicelli, R. C. et al. Synaptic pruning by microglia is necessary for normal brain development. science 333, 1456–1458 (2011).

18 Schafer, D. P. et al. Microglia sculpt postnatal neural circuits in an activity and complement-dependent manner. Neuron 74, 691–705 (2012).

19 Hong, S. et al. Complement and microglia mediate early synapse loss in Alzheimer mouse models. Science 352, 712–716 (2016).

20 Mondo, E. et al. A developmental analysis of juxtavascular microglia dynamics and interactions with the vasculature. Journal of Neuroscience 40, 6503–6521 (2020).

21 Utz, S. G. et al. Early fate defines microglia and non-parenchymal brain macrophage development. Cell 181, 557–573. e518 (2020).

22 Lou, N. et al. Purinergic receptor P2RY12-dependent microglial closure of the injured blood– brain barrier. Proceedings of the National Academy of Sciences 113, 1074–1079 (2016).

23 Haruwaka, K. et al. Dual microglia effects on blood brain barrier permeability induced by systemic inflammation. Nature communications 10, 5816 (2019).

24 Elmore, M. R. et al. Colony-stimulating factor 1 receptor signaling is necessary for microglia viability, unmasking a microglia progenitor cell in the adult brain. Neuron 82, 380–397 (2014).

25 Yi, M.-H. et al. Chemogenetic manipulation of microglia inhibits neuroinflammation and neuropathic pain in mice. Brain, behavior, and immunity 92, 78–89 (2021).

26 Rojo, R. et al. Deletion of a Csf1r enhancer selectively impacts CSF1R expression and development of tissue macrophage populations. Nature communications 10, 3215 (2019).

27 Cox, S. B., Woolsey, T. A. & Rovainen, C. M. Localized dynamic changes in cortical blood flow with whisker stimulation corresponds to matched vascular and neuronal architecture of rat barrels. Journal of Cerebral Blood Flow & Metabolism 13, 899–913 (1993).

28 Sun, Y.-Y. et al. Prophylactic edaravone prevents transient hypoxic-ischemic brain injury: implications for perioperative neuroprotection. Stroke 46, 1947–1955 (2015).

29 Ma, X. et al. Depletion of microglia in developing cortical circuits reveals its critical role in glutamatergic synapse development, functional connectivity, and critical period plasticity. Journal of Neuroscience Research 98, 1968–1986 (2020).

30 Henry, R. J. et al. Microglial depletion with CSF1R inhibitor during chronic phase of experimental traumatic brain injury reduces neurodegeneration and neurological deficits. Journal of Neuroscience 40, 2960–2974 (2020).

31 Lund, H. et al. Competitive repopulation of an empty microglial niche yields functionally distinct subsets of microglia-like cells. Nature communications 9, 1–13 (2018).

32 Klawonn, A. M. et al. Microglial activation elicits a negative affective state through prostaglandin-mediated modulation of striatal neurons. Immunity 54, 225–234. e226 (2021).

33 Thompson, K. J. et al. DREADD agonist 21 is an effective agonist for muscarinic-based DREADDs in vitro and in vivo. ACS Pharmacology & Translational Science 1, 61–72 (2018).

34 Attwell, D. et al. Glial and neuronal control of brain blood flow. Nature 468, 232–243 (2010).

35 Niwa, K., Haensel, C., Ross, M. E. & Iadecola, C. Cyclooxygenase-1 participates in selected vasodilator responses of the cerebral circulation. Circulation research 88, 600–608 (2001).

36 Takano, T. et al. Astrocyte-mediated control of cerebral blood flow. Nature Neuroscience 9, 260–267 (2006).

37 Hahn, T. et al. Neurovascular coupling in the human visual cortex is modulated by cyclooxygenase-1 (COX-1) gene variant. Cerebral cortex 21, 1659–1666 (2011).

38 Butovsky, O. et al. Identification of a unique TGF-β–dependent molecular and functional signature in microglia. Nature Neuroscience 17, 131–143 (2014).

39 Yamada, M. et al. Cholinergic dilation of cerebral blood vessels is abolished in M5 muscarinic acetylcholine receptor knockout mice. Proceedings of the National Academy of Sciences 98, 14096–14101 (2001).

40 Yang, C. et al. Therapeutic Benefits of Adropin in Aged Mice After Transient Ischemic Stroke via Reduction of Blood-Brain Barrier Damage. Stroke (2022).

41 McNamara, N. B. et al. Microglia regulate central nervous system myelin growth and integrity. Nature 613, 120–129 (2023).

42 McKinsey, G. L. et al. A new genetic strategy for targeting microglia in development and disease. elife 9, e54590 (2020).

43 Kim, J.-S. et al. A binary Cre transgenic approach dissects microglia and CNS border-associated macrophages. Immunity 54, 176–190. e177 (2021).

44 Eyo, U. B. et al. Neuronal hyperactivity recruits microglial processes via neuronal NMDA receptors and microglial P2Y12 receptors after status epilepticus. Journal of Neuroscience 34, 10528–10540 (2014).

45 Borgus, J. R., Puthongkham, P. & Venton, B. J. Complex sex and estrous cycle differences in spontaneous transient adenosine. Journal of neurochemistry 153, 216–229 (2020).

46 Santisteban, M. M. et al. Meningeal interleukin-17-producing T cells mediate cognitive impairment in a mouse model of salt-sensitive hypertension. Nature Neuroscience 27, 63–77 (2024).

47 Drieu, A. et al. Parenchymal border macrophages regulate the flow dynamics of the cerebrospinal fluid. Nature 611, 585–593 (2022).

48 Burnstock, G. Historical review: ATP as a neurotransmitter. Trends in pharmacological sciences 27, 166–176 (2006).

49 Fields, R. D. & Burnstock, G. Purinergic signalling in neuron–glia interactions. Nature Reviews Neuroscience 7, 423–436 (2006).

50 Wells, J. A. et al. A critical role for purinergic signalling in the mechanisms underlying generation of BOLD fMRI responses. Journal of Neuroscience 35, 5284–5292 (2015).

51 Kitajima, N. et al. Real-time in vivo imaging of extracellular ATP in the brain with a hybrid-type fluorescent sensor. Elife 9, e57544 (2020).

52 Faroqi, A. H. et al. In vivo detection of extracellular adenosine triphosphate in a mouse model of traumatic brain injury. Journal of neurotrauma 38, 655–664 (2021).

53 Wang, X. et al. P2X7 receptor inhibition improves recovery after spinal cord injury. Nature medicine 10, 821–827 (2004).

54 Cai, C. et al. Stimulation-induced increases in cerebral blood flow and local capillary vasoconstriction depend on conducted vascular responses. Proceedings of the National Academy of Sciences 115, E5796–E5804 (2018).

55 Szalay, G. et al. Microglia protect against brain injury and their selective elimination dysregulates neuronal network activity after stroke. Nature communications 7, 11499 (2016).

56 Williams, M. Adenosine: the prototypic neuromodulator. Neurochemistry international 14, 249–264 (1989).

57 Phillis, J. W., Kostopoulos, G. K. & Limacher, J. J. A potent depressant action of adenine derivatives on cerebral cortical neurones. European Journal of Pharmacology 30, 125–129 (1975).

58 Manzoni, O. J., Manabe, T. & Nicoll, R. A. Release of adenosine by activation of NMDA receptors in the hippocampus. Science 265, 2098–2101 (1994).

59 Merlini, M. et al. Microglial Gi-dependent dynamics regulate brain network hyperexcitability. Nature Neuroscience 24, 19–23 (2021).

60 Korte, N., Nortley, R. & Attwell, D. Cerebral blood flow decrease as an early pathological mechanism in Alzheimer’s disease. Acta neuropathologica 140, 793–810 (2020).

61 Pietrowski, M. J. et al. Glial purinergic signaling in neurodegeneration. Frontiers in neurology 12, 654850 (2021).

62 Keren-Shaul, H. et al. A unique microglia type associated with restricting development of Alzheimer’s disease. Cell 169, 1276–1290. e1217 (2017).

63 Butovsky, O. et al. Targeting mi R-155 restores abnormal microglia and attenuates disease in SOD 1 mice. Annals of neurology 77, 75–99 (2015).

## Bibliography for Methods

1 Rothweiler, S. et al. Selective deletion of ENTPD1/CD39 in macrophages exacerbates biliary fibrosis in a mouse model of sclerosing cholangitis. Purinergic signalling 15, 375–385 (2019).

2 Badimon, A. et al. Negative feedback control of neuronal activity by microglia. Nature 586, 417–423 (2020).

3 Takenaka, M. C. et al. Control of tumor-associated macrophages and T cells in glioblastoma via AHR and CD39. Nature Neuroscience 22, 729–740 (2019).

4. Chen, H.-R., et al. Creatine transporter deficiency impairs stress adaptation and brain energetics homeostasis. JCI insight 6 (2021).

5 Sun, Y.-Y. et al. Prophylactic edaravone prevents transient hypoxic-ischemic brain injury: implications for perioperative neuroprotection. Stroke 46, 1947–1955 (2015).

6 Faraco, G. et al. Dietary salt promotes cognitive impairment through tau phosphorylation. Nature 574, 686–690 (2019).

7 Park, L. et al. Tau induces PSD95–neuronal NOS uncoupling and neurovascular dysfunction independent of neurodegeneration. Nature Neuroscience 23, 1079–1089 (2020).

8 Bisht, K. et al. Capillary-associated microglia regulate vascular structure and function through PANX1-P2RY12 coupling in mice. Nature communications 12, 1–13 (2021).

9 Chen, H.-R. et al. Monocytes promote acute neuroinflammation and become pathological microglia in neonatal hypoxic-ischemic brain injury. Theranostics 12, 512 (2022).

10 Ewels, P., Magnusson, M., Lundin, S. & Käller, M. MultiQC: summarize analysis results for multiple tools and samples in a single report. Bioinformatics 32, 3047–3048 (2016).

11 Dobin, A. et al. STAR: ultrafast universal RNA-seq aligner. Bioinformatics 29, 15–21 (2013).

12 Anders, S., Pyl, P. T. & Huber, W. HTSeq—a Python framework to work with high-throughput sequencing data. Bioinformatics 31, 166–169 (2015).

13 Love, M. I., Huber, W. & Anders, S. Moderated estimation of fold change and dispersion for RNA-seq data with DESeq2. Genome biology 15, 1–21 (2014).

14 Subramanian, A. et al. Gene set enrichment analysis: a knowledge-based approach for interpreting genome-wide expression profiles. Proceedings of the National Academy of Sciences 102, 15545–15550 (2005).

15 Swamy, B. K. & Venton, B. J. Subsecond detection of physiological adenosine concentrations using fast-scan cyclic voltammetry. Analytical chemistry 79, 744–750 (2007).

16 Ganesana, M. & Venton, B. J. Spontaneous, transient adenosine release is not enhanced in the CA1 region of hippocampus during severe ischemia models. Journal of Neurochemistry 159, 887–900 (2021).

17 Ganesana, M. & Venton, B. J. Early changes in transient adenosine during cerebral ischemia and reperfusion injury. PloS one 13, e0196932 (2018).

