## Supplemental Figures for "Microglia modulate cerebral blood flow and neurovascular coupling through ectonucleotidase CD39"

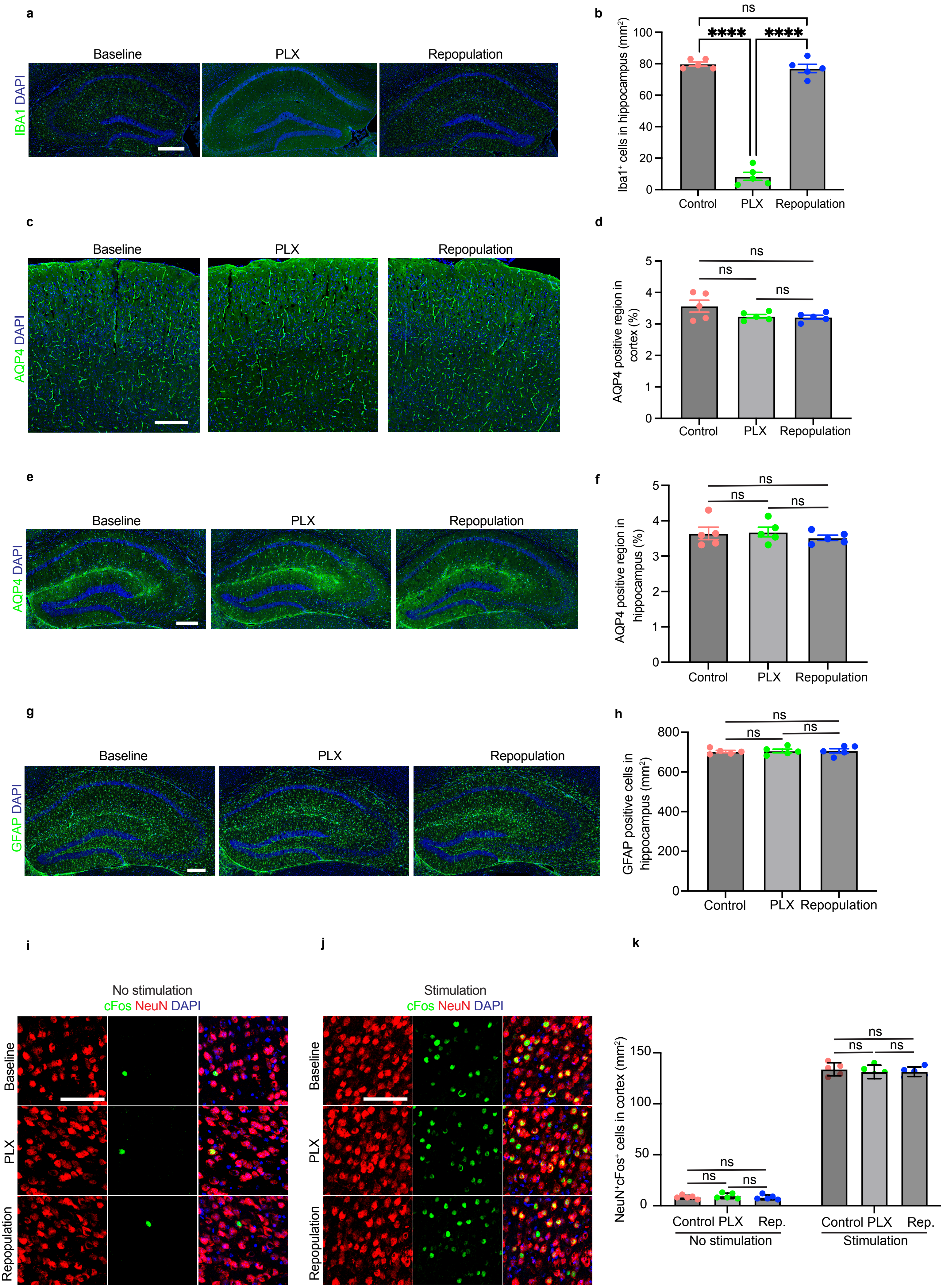

a
Baseline
PLX
Repopulation
IBA1 DAPI
b
ns
****
****
Iba1+ cells in hippocampus (mm2)
100
80
60
40
20
0
Control
PLX
Repopulation
c
Baseline
PLX
Repopulation
AQP4 DAPI
d
ns
ns
ns
AQP4 positive region in cortex (%)
5
4
3
2
1
0
Control
PLX
Repopulation
e
Baseline
PLX
Repopulation
AQP4 DAPI
f
ns
ns
ns
AQP4 positive region in hippocampus (%)
5
4
3
2
1
0
Control
PLX
Repopulation
g
Baseline
PLX
Repopulation
GFAP DAPI
h
ns
ns
ns
GFAP positive cells in hippocampus (mm2)
800
600
400
200
0
Control
PLX
Repopulation
i
No stimulation
cFos NeuN DAPI
Baseline
PLX
Repopulation
j
Stimulation
cFos NeuN DAPI
Baseline
PLX
Repopulation
k
NeuN+cFos+ cells in cortex (mm2)
150
100
50
0
ns
ns
ns
ns
ns
ns
Control PLX Rep.
No stimulation
Control PLX Rep.
Stimulation

**Extended Data Figure 1. Microglia depletion or repopulation did not induce astrocyte activation or change whisker stimulation induced neuronal activation. a,** Immuno-histochemistry images showing $IBA1^+$ cells co-stained with DAPI in baseline, PLX and repopulation conditions in hippocampus. Bar=400μm. **b,** Quantification of $IBA1^+$ cells in hippocampus in three conditions (baseline, PLX and repopulation). Data are presented as mean ± s.e.m. p-values were determined by one-way ANOVA with Tukey post hoc test. **c-f,** Immunohistochemistry images showing AQP4 expression (co-stained with DAPI) in cortex (**c**) and hippocampus (**e**) at baseline, PLX and repopulation conditions. Bar=300μm. Quantification of AQP4 expression in cortex (**d**) and hippocampus (**f**) in three conditions (baseline, PLX and repopulation). Data are presented as mean ± s.e.m. p-values were determined by one-way ANOVA with Tukey post hoc test. **g,** Immunohistochemistry images showing GFAP expression (co-stained with DAPI) in hippocampus at baseline, PLX and repopulation conditions. Bar=300μm. **h,** Quantification of GFAP expression in hippocampus in three conditions (baseline, PLX and repopulation). Data are presented as mean ± s.e.m. p-values were determined by one-way ANOVA with Tukey post hoc test. **i-j,** Representative images showing cFos, NeuN, DAPI staining in barrel cortex without whisker stimulation (**i**) or with whisker stimulation (**j**) in three conditions. Bar=400μm. **k,** Quantification of cFos expression in barrel cortex with or without whisker stimulation in three conditions (baseline, PLX and repopulation). Data are presented as mean ± s.e.m. p-values were determined by one-way ANOVA with Tukey post hoc test.

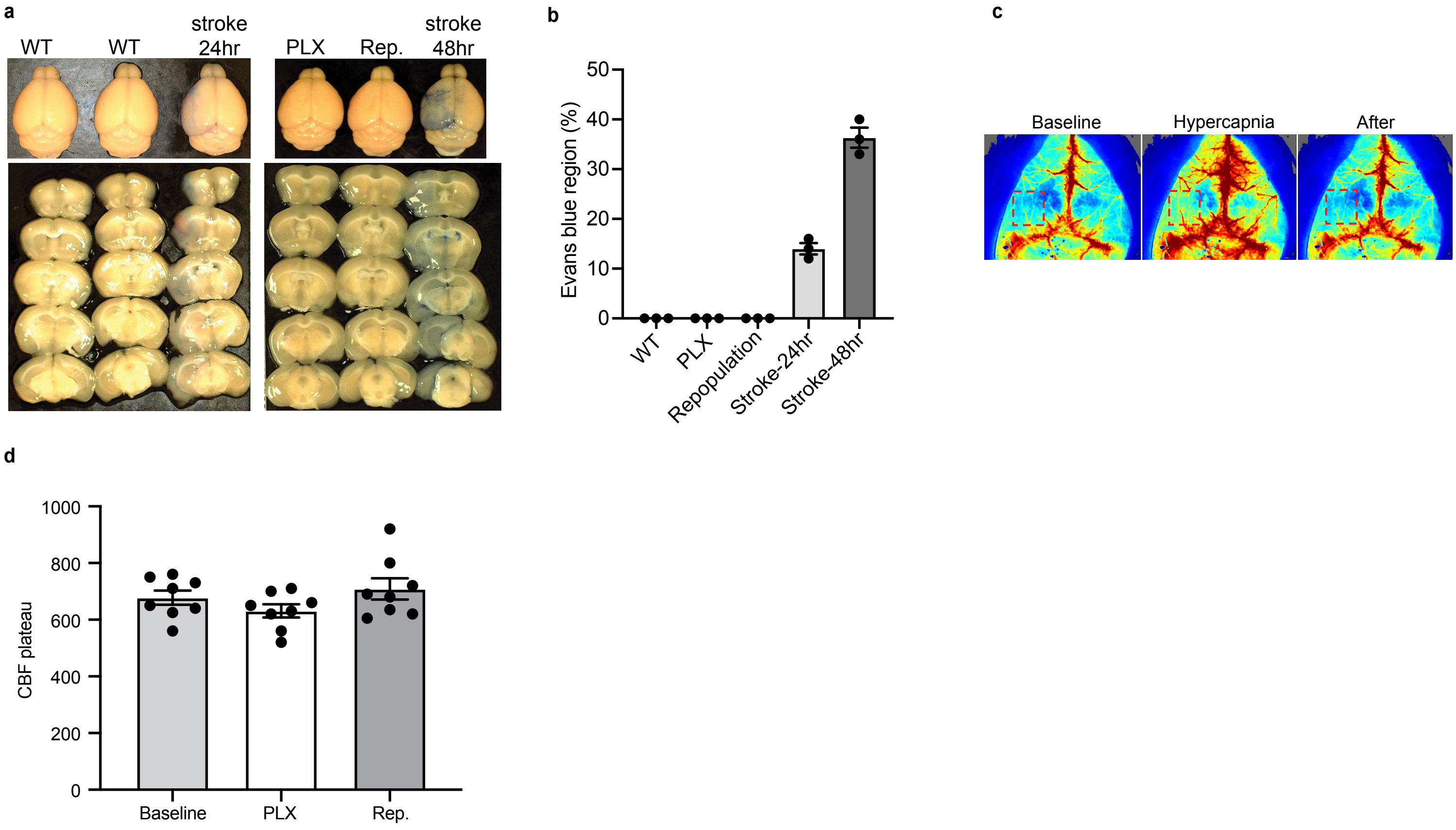
a
WT
WT
stroke 24hr
PLX
Rep.
stroke 48hr
b
Evans blue region (%)
50
40
30
20
10
0
WT
PLX
Repopulation
Stroke-24hr
Stroke-48hr
c
Baseline
Hypercapnia
After
d
CBF plateau
1000
800
600
400
200
0
Baseline
PLX
Rep.

**Extended Data Figure 2. Microglia depletion or repopulation did not change blood brain barrier permeability. a,** Mice were injected with Evans Blue (EB) (2%, 2 ml/kg) from tail vein 1.5 h before perfusion. Representative images showing EB leakage into brain parenchyma in different conditions. **b,** Quantification of blood brain barrier leakage with EB in different conditions. **c,** Representative images of hypercapnia (8% $CO_2$) induced CBF changes. Red dash line square marked interested regions. **d,** CBF plateau value during hypercapnia stimulation at three different time points.

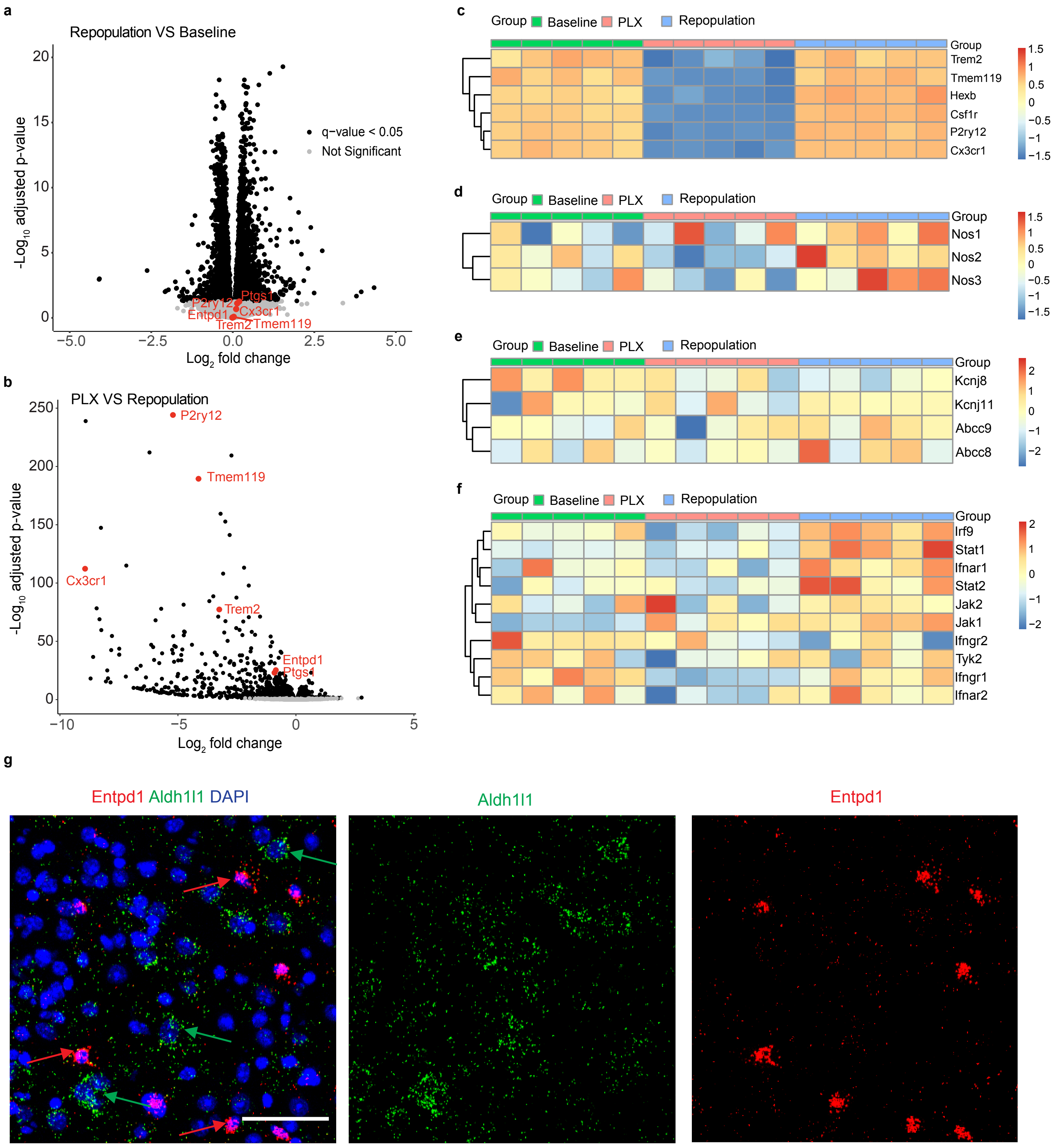

a
Repopulation VS Baseline
-Log10 adjusted p-value
Log2 fold change
q-value < 0.05
Not Significant
P2ry12
Ptgs1
Entpd1
Cx3cr1
Trem2
Tmem119
b
PLX VS Repopulation
-Log10 adjusted p-value
Log2 fold change
P2ry12
Tmem119
Cx3cr1
Trem2
Entpd1
Ptgs1
c
Group
Baseline
PLX
Repopulation
Trem2
Tmem119
Hexb
Csf1r
P2ry12
Cx3cr1
d
Nos1
Nos2
Nos3
e
Kcnj8
Kcnj11
Abcc9
Abcc8
f
Irf9
Stat1
Ifnar1
Stat2
Jak2
Jak1
Ifngr2
Tyk2
Ifngr1
Ifnar2
g
Entpd1 Aldh1l1 DAPI
Aldh1l1
Entpd1

**Extended Data Figure 3. Bulk RNAseq analysis showing gene expression changes in baseline, PLX3392 treated and microglia repopulation mice and CD73 expression in different brain regions. a,** Volcano plot of the cortex genes showing the magnitude (log2 [fold change]) and probability (−log10 [adjusted p-value]) in microglia repopulation group versus baseline group. Black and gray dots represent significantly changed and non-significantly changed genes, respectively. **b,** Volcano plot of the cortex genes showing the magnitude (log2 [fold change]) and probability (−log10 [adjusted p-value]) in PLX3392-treated group versus repopulation group. Black and gray dots represent significantly changed and non-significantly changed genes, respectively. **c,** Heat map showing microglia marker genes' expression in 5 samples from each group (baseline, PLX, Repopulation) in row z scores of log2 CPM values. **d,** Heat map showing nitric oxide synthase (NOS) gene expression in five samples of each group in row z scores of log2 CPM values. **e,** Heat map showing ATP sensitive potassium channel gene expression in five samples of each group in row z scores of log2 CPM values. **f,** Heat map showing interferon related gene expression in five samples of each group in row z scores of log2 CPM values. **g,** Representative images of multi-color RNAscope experiments showing CD39 (Entpd1, red color) and astrocyte marker gene Aldh1l1 expression in brain cortex. CD39 did not co-localize with Aldh1l1. Bar=50μm.

a

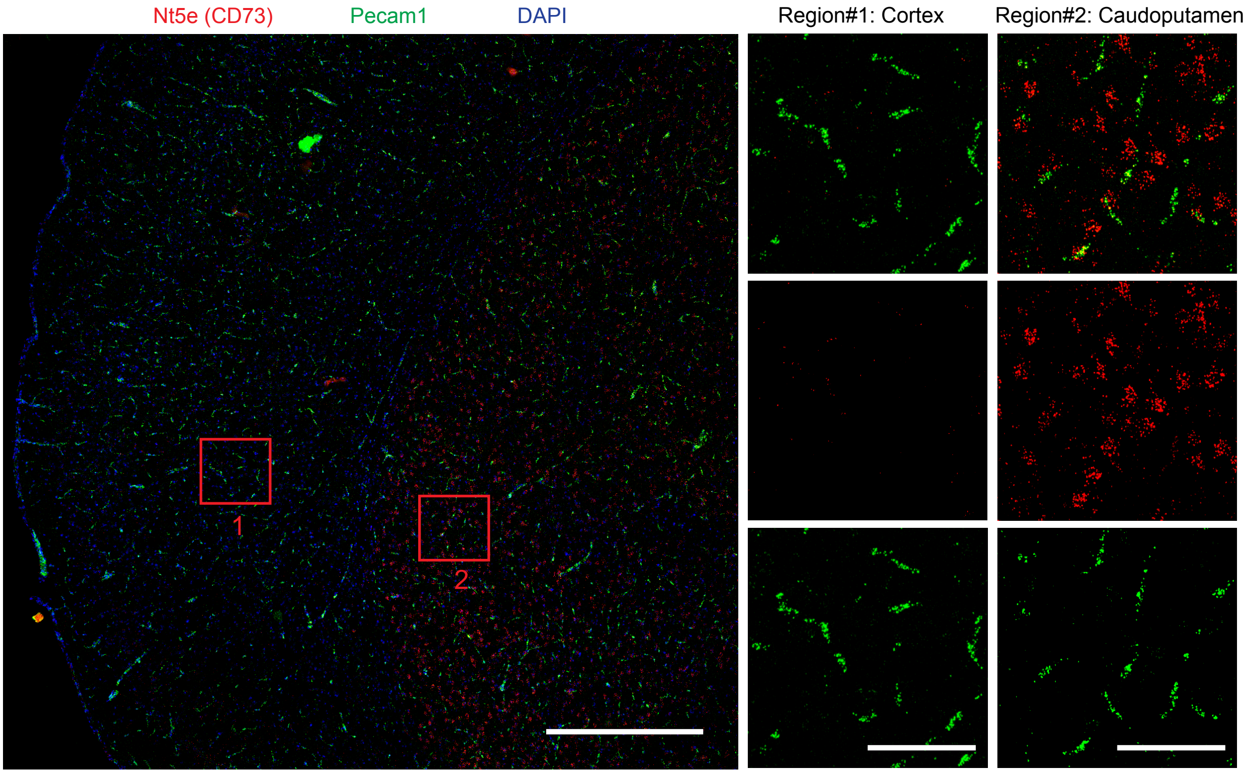

Nt5e (CD73)
Pecam1
DAPI
Region#1: Cortex
Region#2: Caudoputamen
1
2


b

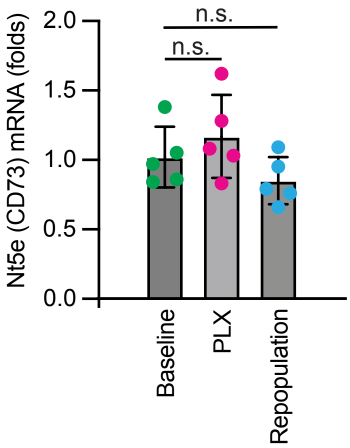

Nt5e (CD73) mRNA (folds)
2.0
1.5
1.0
0.5
0.0
n.s.
n.s.
Baseline
PLX
Repopulation

**Extended Data Figure 4. In situ hybridization results showing CD73 (Nt5e) differential expression in cortex and caudoputamen. a,** Representative images of multi-color RNAscope experiments showing CD73 (Nt5e, red color) expression in different brain regions (co-stained with endothelial cells marker gene Pecam1, green color). CD73 expression is low in cortex (region #1) and high in caudoputamen (region #2). Left panel, bar=500μm; middle and right panel, bar=100μm. **b,** RT-qPCR verification of CD73 (Nt5e) expression changes in three conditions: baseline, PLX3397-treated and repopulation. Data are mean ± s.e.m. Each dot represents one mouse. p-values were determined by one-way ANOVA with Tukey post hoc test.

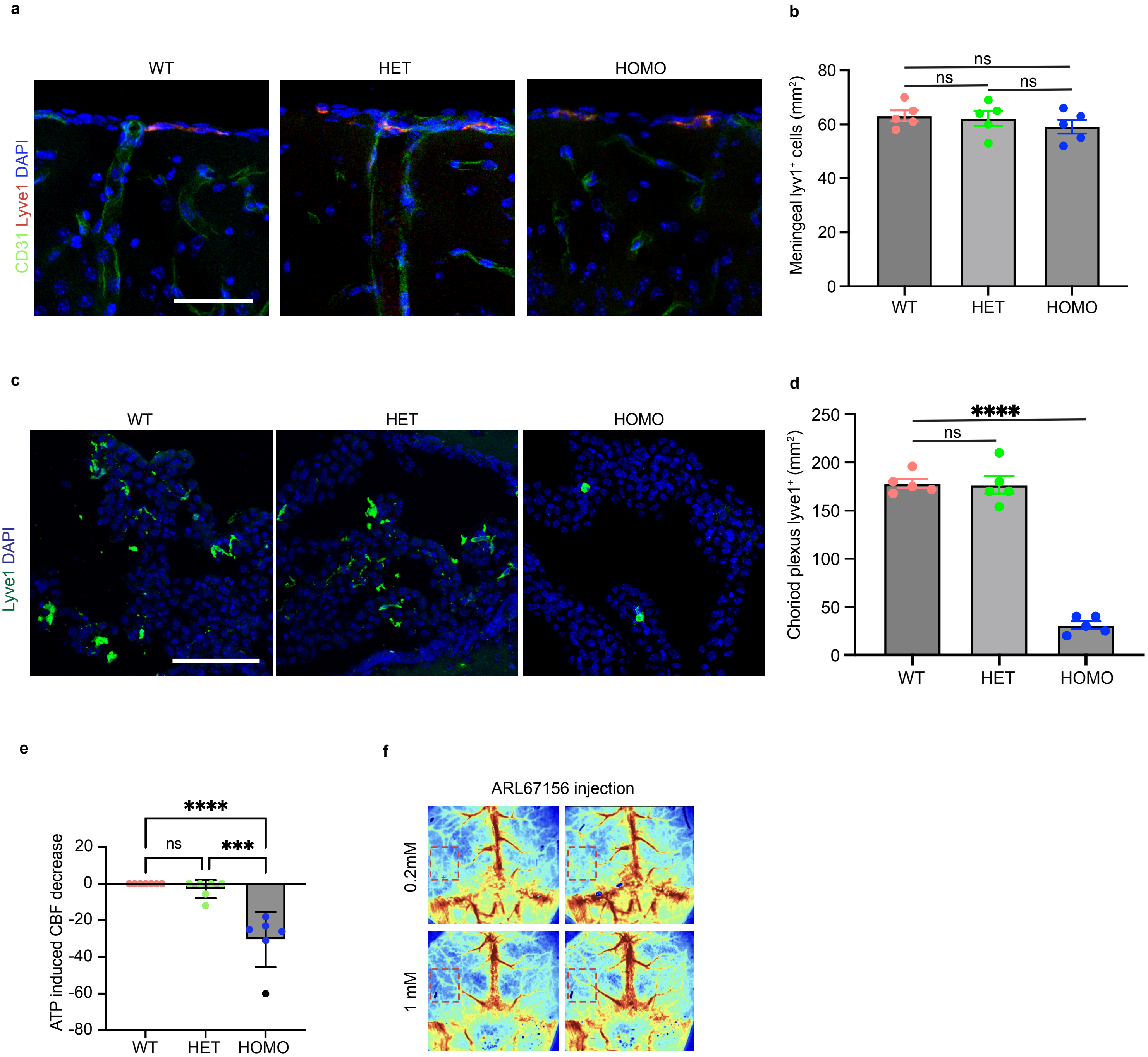
a
WT
HET
HOMO
CD31 Lyve1 DAPI
b
Meningeal lyv1⁺ cells (mm²)
ns
ns
ns
0
20
40
60
80
WT
HET
HOMO
c
WT
HET
HOMO
Lyve1 DAPI
d
Choriod plexus lyve1⁺ (mm²)
****
ns
0
50
100
150
200
250
WT
HET
HOMO
e
ATP induced CBF decrease
****
ns
***
20
0
-20
-40
-60
-80
WT
HET
HOMO
f
ARL67156 injection
0.2mM
1 mM

**Extended Data Figure 5. Characterization of brain border associated macrophages (BAMs) in Csf1r$^{\Delta FIRE/\Delta FIRE}$ mice. a,** Representative immunohistochemistry images showing lyve1$^{+}$ macrophages in leptomeninges of WT, heterozygous and homozygous Csf1r$^{\Delta FIRE/\Delta FIRE}$ mice. Bar= 200μm. **b,** Quantification of lyve1$^{+}$ macrophages in leptomeninges of WT, heterozygous and homozygous Csf1r$^{\Delta FIRE/\Delta FIRE}$ mice. Data are presented as mean ± s.e.m. p-values were determined by one-way ANOVA with Tukey post hoc test. **c,** Representative immunohistochemistry images showing lyve1$^{+}$ macrophages in choroid plexus of WT, heterozygous and homozygous Csf1r$^{\Delta FIRE/\Delta FIRE}$ mice. Bar= 100μm. **d,** Quantification of lyve1$^{+}$ macrophages in choroid plexus of WT, heterozygous and homozygous Csf1r$^{\Delta FIRE/\Delta FIRE}$ mice. Data are presented as mean ± s.e.m. p-values were determined by one-way ANOVA with Tukey post hoc test. **e,** Quantification of ATP ICM injection induced CBF decrease in early phase of injection in WT, HET, and HOMO of Csf1r$^{\Delta FIRE/\Delta FIRE}$ mice. p-values were determined by one-way ANOVA with Tukey post hoc test. Data are presented as mean ± s.e.m. **f,** Representative micrographs showing ICM injection of CD39 inhibitor ARL67156 induced CBF change at different concentrations.

**Entpd1 fl/fl**

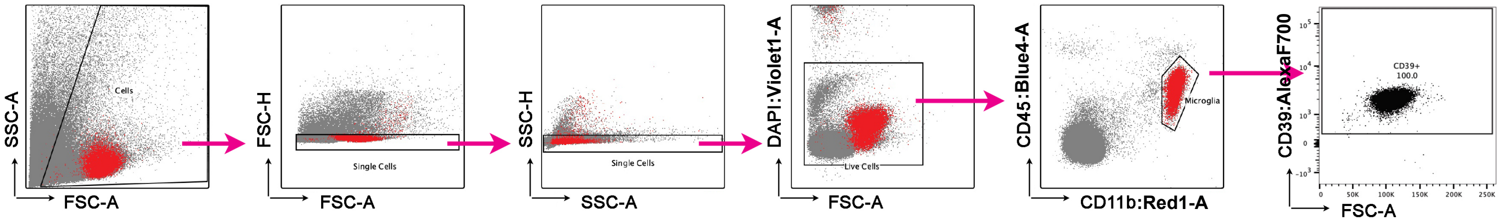


**CX3CR1-CreERT2; Entpd1 fl/fl**

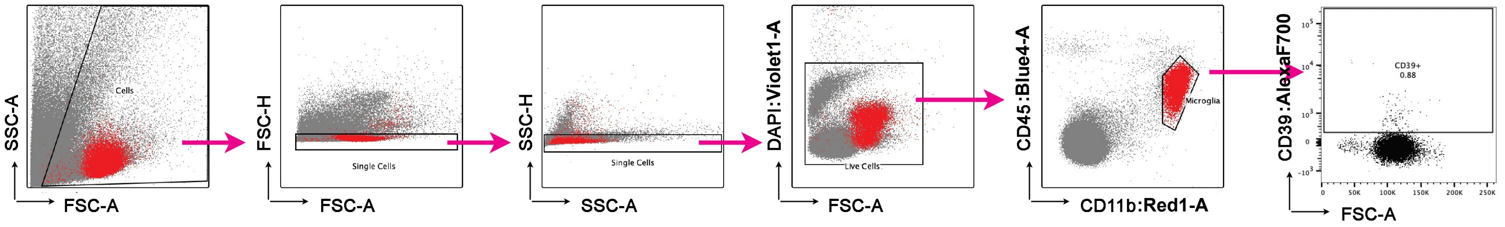

**Extended Data Figure 6. Flow cytometry data showing efficient microglia CD39 conditional knockout in Cx3cr1-CreER-;Entpd1 (CD39)fl/fl after tamoxifen injection.** Surface expression of CD39 in isolated cortical cells from adult Entpd1fl/fl mice (top row) or Cx3cr1-CreER-;Entpd1 (CD39)fl/fl littermates (bottom row) mice by fluorescence-activated cell sorting. Doublets were removed based on FSC and SSC, then dead cells were removed based on DAPI positivity. Stained Entpd1fl/fl and Cx3cr1-CreER-;Entpd1 (CD39)fl/fl samples were sorted for CD11bhigh and CD45low expression to identify microglia. CD39 surface expression is shown in the last panel of each row with their respective population percentages. Back gating is shown in red.

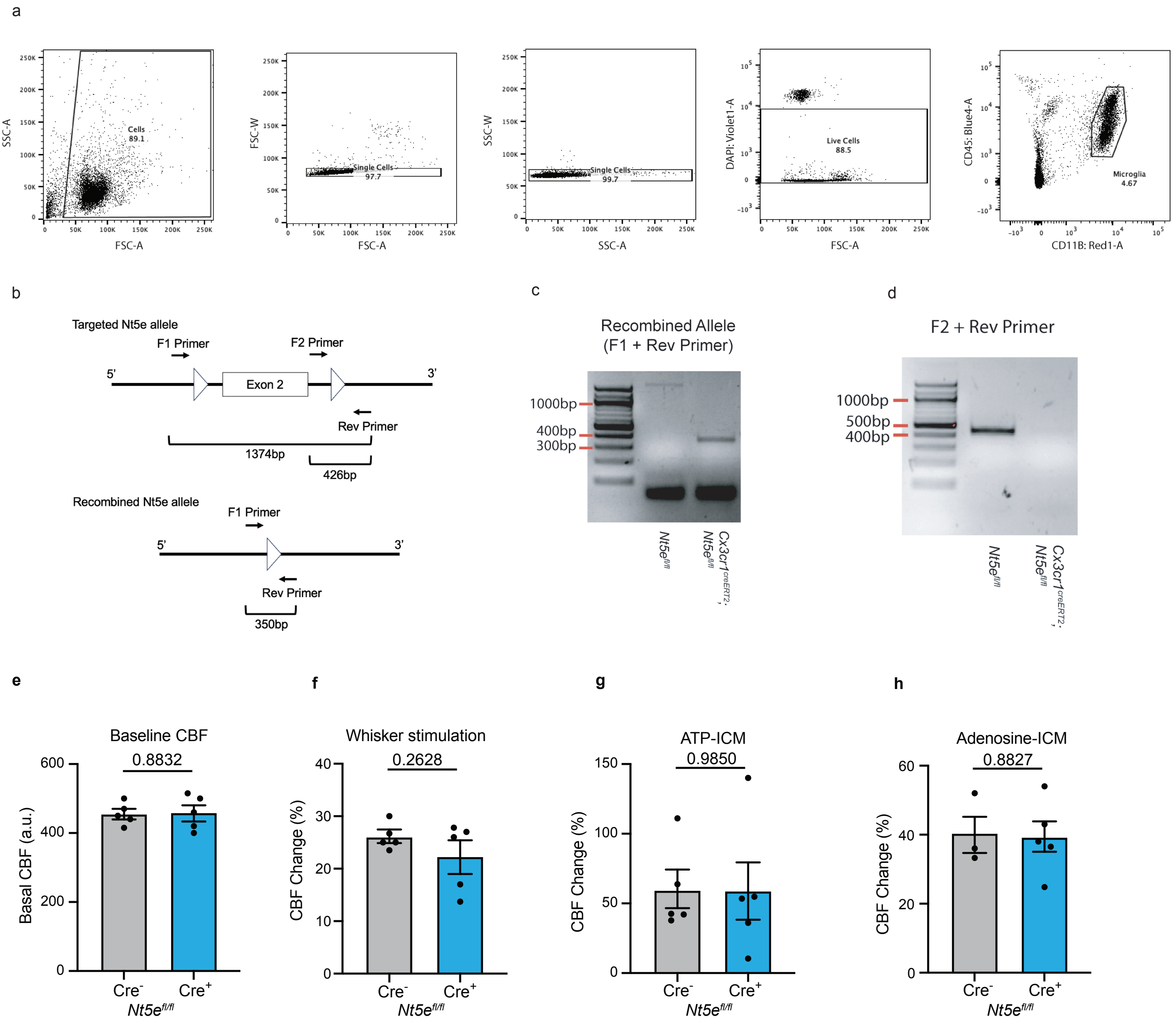

a
SSC-A
FSC-A
Cells
89.1
FSC-W
Single Cells
97.7
SSC-W
SSC-A
Single Cells
99.7
DAPI: Violet1-A
Live Cells
88.5
CD45: Blue4-A
CD11B: Red1-A
Microglia
4.67
b
Targeted Nt5e allele
F1 Primer
F2 Primer
5'
Exon 2
3'
Rev Primer
1374bp
426bp
Recombined Nt5e allele
F1 Primer
5'
3'
Rev Primer
350bp
c
Recombined Allele
(F1 + Rev Primer)
1000bp
400bp
300bp
Nt5e fl/fl
Cx3cr1 creERT2; Nt5e fl/fl
d
F2 + Rev Primer
1000bp
500bp
400bp
Nt5e fl/fl
Cx3cr1 creERT2; Nt5e fl/fl
e
Baseline CBF
0.8832
Basal CBF (a.u.)
Cre-
Cre+
Nt5e fl/fl
f
Whisker stimulation
0.2628
CBF Change (%)
Cre-
Cre+
Nt5e fl/fl
g
ATP-ICM
0.9850
CBF Change (%)
Cre-
Cre+
Nt5e fl/fl
h
Adenosine-ICM
0.8827
CBF Change (%)
Cre-
Cre+
Nt5e fl/fl

**Extended Data Figure 7. CD73 (Nt5e) was low expressed in cortex region and microglial CD73 conditional knockout did not change neurovascular coupling in barrel cortex. a,** Representative images of multi-color RNAscope experiments showing CD73 (Nt5e, red color) expression in different brain regions (co-stained with endothelial cells marker gene Pecam1, green color). CD73 expression is low in cortex (region #1) and high in caudoputamen (region #2). Left panel, bar=500μm; middle and right panel, bar=100μm. **b-d,** RT-qPCR verification of CD73 (Nt5e) expression changes in three conditions: baseline, PLX3397-treated and repopulation. Data are mean ± s.e.m. Each dot represents one mouse. p-values were determined by one-way ANOVA with Tukey post hoc test. **e-h,** Comparison of the baseline CBF (**e**), CBF change to whisker stimulation (**f**), CBF change to ICM-ATP injection (**g**), and CBF change to ICM-adenosine injection (**h**) in tamoxifen-induced *Cx3cr1-CreER*$^{-}$*;Nt5e (CD73)*$^{fl/fl}$ versus *Cx3cr1-CreER*$^{+}$*;Nt5e (CD73)*$^{fl/fl}$ mice. Note that microglial CD73-deletion failed to significantly change the baseline CBF (**e**) or CBF change to whisker stimulation (**f**), ICM-ATP injection (**g**) or ICM-adenosine injection (**h**). Data are presented as mean ± s.e.m. p-values were determined by unpaired *t* test.
